# Detecting DNA methylation patterns suggestive of variable escape from X-chromosome inactivation

**DOI:** 10.64898/2026.06.15.732395

**Authors:** Qicheng Zhao, Ohanna C. L. Bezerra, Kathleen Oros Klein, Mark Lamin, Marie-Claude Beaulieu, Marc Rodger, Michael J. Kovacs, Liam O’Neil, Carolyn Brown, Marie Hudson, Ines Colmegna, Sasha Bernatsky, France Gagnon, Anna K. Naumova, Qihuang Zhang, Celia M. T. Greenwood

**Author notes:** **Corresponding author:** Celia M. T. Greenwood, Lady Davis Institute, Sir Mortimer B. Davis Jewish General Hospital CIUSSS du Centre-Ouest-de-l’Île-de-Montréal, Montreal, QC, Canada H3T 1A2. Co-first authors (Qicheng Zhao, Ohanna C. L. Bezerra).

## Abstract

The X chromosome is often excluded from studies analyzing associations between traits and DNA methylation. In females, one copy of most genes on the X is inactivated (X-chromosome inactivation; XCI) through DNA methylation of the gene promoter on the inactive X. This leads to challenges in analyzing and interpreting DNA methylation data patterns. Particularly for sex-biased diseases and traits, there may be many loci of interest on the X chromosome, which contains about 5% of the genome. To address the need for appropriate analysis of DNA methylation data on the X chromosome, we develop a statistical approach to infer locus-specific escape from XCI sensitive to phenotype or covariate values. Performance of this method is illustrated by analysis of data from two sex-biased traits: rheumatoid arthritis which is 3-fold more common in females, and recurrent venous thromboembolism which occurs 2.5 times more often in males. Analyses of these two datasets identify new trait-associated loci on the X chromosome, demonstrate the capabilities of the new method for both bisulfite sequencing data and Illumina EPIC data, suggest at least one locus where variable escape may explain a sex-specific disease association, and rule out variable escape as a potential explanation at other loci.

**Graphical abstract:** 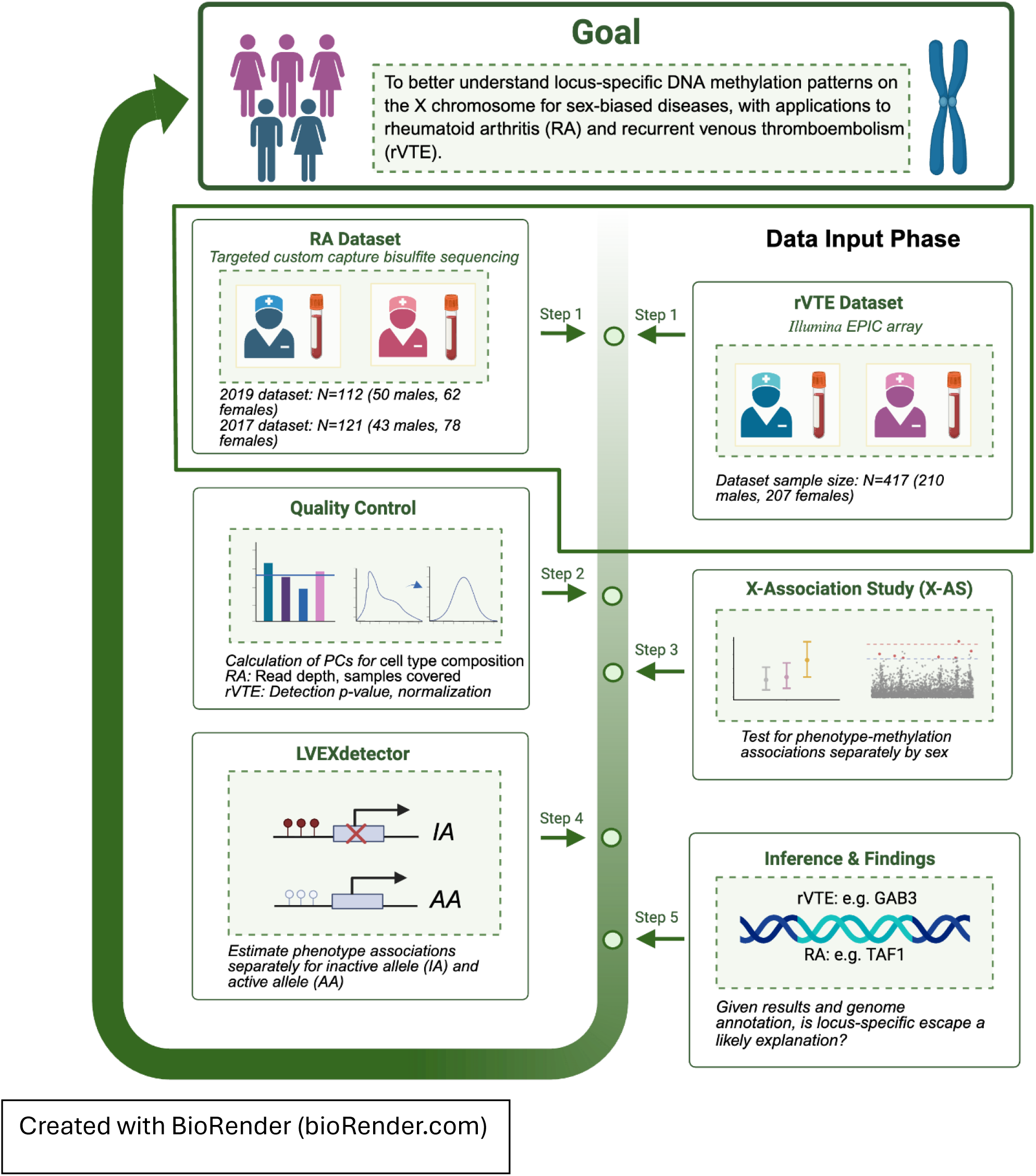

Created with BioRender (bioRender.com)

## 1 Introduction

The X chromosome poses unique challenges and opportunities in genetic and epigenetic association studies. In mammals, females possess two X chromosomes, whereas males carry one X and one Y chromosome. To achieve dosage compensation, one of the two X chromosomes in the somatic cells of females undergoes transcriptional silencing early in embryonic development {Lyon, 1972 #4164;Lyon, 1961 #4050}. The process of X-chromosome inactivation (XCI) is random; that is, one of the two chromosomes in each cell is inactivated (*Xi*), leading to a cellular mosaicism with respect to expression and epigenetic markers, i.e. either chromosome could be the active X (*Xa*). However, certain X-linked genes escape X-inactivation. These include genes in the pseudo-autosomal regions that have homologues on the Y chromosome and hence do not require dosage compensation, and additional genes that are dispersed among those that are subject to XCI (Balaton and Brown 2016). Such genes remain transcriptionally active on both X chromosomes, albeit expression from the *Xi* may not reach the levels of the *Xa* (Balaton and Brown 2016; Carrel and Brown 2017; Brown et al. 1991; Penny et al. 1996). Furthermore, there can be variability in the escape status in different tissues (Berletch et al. 2015) as well as between individuals (Balaton and Brown 2016). For example, ageing may lead to escape from XCI at certain loci (Roberts et al. 2022).

Variation in XCI patterns, arising from mutations in X-inactivation specific transcript (*XIST*) or locus-specific variable escape from XCI, has been proposed as an explanation for sex biases in disease rates (Lovell and Anguera 2025; Zhang et al. 2020; Miquel, Faz-Lopez, and Guery 2023; Huret et al. 2024). Although methodology for detecting associations between disease and genetic variants on the X chromosome has existed for some time (Derkach, Lawless, and Sun 2014; Deng et al. 2019), methods applicable to epigenetic variation remain underdeveloped, leading to the untapped potential of existing chromosome X epigenetic data, such as DNA methylation from epigenome-wide association studies (EWAS).

DNA methylation is widely implicated in the regulation of gene expression, genome architecture and stability. It is also an important marker of XCI (Cotton et al. 2011). Variation in DNA methylation is involved in the pathogenesis of autoimmune disorders, cancer, and other common human diseases. Variable escape is proposed to contribute to the observed sex bias in several autoimmune diseases (Huret et al. 2024; Jin and Liu 2018; Robertson 2005; Richardson 2003). Exclusion of the X chromosome from DNA methylation association studies arises since a naïve statistical analysis would be inappropriate due to XCI, and reflects the scarcity of methods to detect and interpret altered epigenetic patterns in the presence of XCI (Inkster et al. 2023).

In very simple terms, when a gene’s promoter region is highly methylated, the gene is likely silent (repressed). In fact, promoters of actively transcribed genes that are subject to XCI are methylated on *Xi* and unmethylated on *Xa*, whereas promoters of actively transcribed genes that escape XCI are unmethylated on both *Xi* and *Xa* (Cotton et al. 2011; Cotton et al. 2015). Therefore, we propose the use of DNA methylation data to statistically infer a locus’ XCI state (escape versus subject to XCI), as well as the methylation levels on *Xi* and *Xa* at that locus. Following from this hypothesis, we posit an analytic pipeline for identifying X-chromosome loci associated with sex-biased diseases or phenotypes, including a new method to estimate phenotype-associated DNA methylation patterns consistent with locus-specific variable escape from XCI. For simplicity, we abbreviate *Locus-specific Variable Escape from XCI* as LVE-X, and this new method can also rule out LVE-X as a potential explanation. To illustrate the performance of the proposed methodology, we explore X-chromosome DNA methylation patterns in studies of two sex-biased conditions, rheumatoid arthritis, which is more common in females, and recurrent venous thromboembolism which is more common in males.

The X chromosome harbors a high density of immune-related genes, and loci on the X chromosome are associated with several auto-immune diseases (Bianchi et al. 2012). Variable escape has been proposed to contribute to the observed sex bias in autoimmune disease (Huret et al. 2024). Rheumatoid arthritis (RA) is a chronic autoimmune arthritis with a marked sex bias in prevalence (three-fold more females than males affected) (Gerosa et al. 2008). Given the associations between escape from XCI and autoimmunity, LVE-X may be a potential contributing factor to the female predominance observed in RA (Lovell and Anguera 2025). We hypothesize that higher expression due to escape from XCI of certain immunity-related X-linked genes in some individuals (LVE-X) leads to increased RA risk.

Conversely, individuals with lower expression at such genes, i.e. males or females with no escape, are at lower risk of developing RA.

Beyond autoimmunity, sex-biased regulation of immune and coagulation pathways is also central to thrombotic disease, providing a broader context in which X-linked regulatory mechanisms may contribute to disease risk. Venous thromboembolism (VTE), including deep vein thrombosis (DVT) and pulmonary embolism (PE), is a multifactorial disease that can be classified as unprovoked, with no identifiable risk factors, or provoked by persistent (e.g., cancer) or transient (e.g., surgery) pro-thrombotic factors. This classification guides recommendations for the duration of anticoagulation treatment, as recurrence risk differs between provoked and unprovoked events. Although rates of a first VTE are similar between sexes, for unprovoked VTE, males have 2.5-fold higher risk of recurrence (rVTE) after anticoagulant discontinuation compared with females (Kyrle et al. 2004). Exogenous female sex hormones, although associated with first VTE, are not independent predictors of recurrent VTE; nor dictate differential VTE management or are known modifiable risk factors (Roach, Cannegieter, and Lijfering 2014; Rodger et al. 2017; Le Gal et al. 2010). Thus, the biological basis of this sex difference remains unclear. Increasing attention focuses on the involvement of X-or Y-linked mutations and changes in regulation of genes exhibiting sex-specific effects (Roach, Cannegieter, and Lijfering 2014). Several components of the coagulation cascade are encoded on the X chromosome, including *F8* and *F9*, for which gene duplications and variants contribute to VTE risk (Simioni et al. 2021; Thibord et al. 2022; Lindstrom et al. 2019). Studies of Y-chromosome variation have not supported a male-specific genetic predisposition to either first or recurrent VTE (de Haan et al. 2016), suggesting that X-linked dosage effects may be more relevant. In a recent study of rVTE using data from the REVERSE I study (Rodger et al. 2008), we identified autosomal genes with pronounced sex differences in DNA methylation; however, whether these reflect treatment effects or intrinsic sex-specific regulation remains unresolved, and warrants replication (Bezerra et al. 2025). Limited research investigated the potential role of X-linked variation in first VTE risk, at the genomic or epigenomic level, and no one has examined it in rVTE.

Evidence suggests that skewed XCI, with predominant silencing of one set of alleles, can protect females from manifesting clinical phenotypes (Sun et al. 2021). We hypothesize that sex-dependent dosage effects arising from locus-specific incomplete XCI could modulate expression of X-linked genes involved in VTE, thereby partially protecting females from rVTE risk.

In this paper, we present a novel method and analytic pipeline for detecting DNA methylation patterns consistent with LVE-X, using two illustrative datasets of well-established sex-biased diseases: RA and rVTE; the two studies used different platforms to measure DNA methylation. We first performed X-chromosome association analyses (X-AS) for each trait separately by sex. After describing the new method to test for LVE-X, we then used it to evaluate whether LVE-X can explain observed sex-specific associations. We report findings relevant to our hypothesis of LVE-X, and novel X-chromosome associations between DNA methylation in males and females, for both RA and rVTE.

## 2 Methods

Using the methods and analytic pipeline developed and described below for detecting DNA methylation patterns consistent or inconsistent with LVE-X, we analyzed two illustrative datasets. The first data set used bisulfite sequencing of stored peripheral blood samples with a custom targeted library, and contains individuals with and without RA. We refer to this as the RA dataset. The second dataset was collected to study rVTE, and measured DNA methylation in blood DNA with the Illumina Methylation EPIC array. We refer to this as the rVTE dataset.

### 2.1 Pipeline and method to estimate LVE-X

Since differentially-methylated loci on the X chromosome may arise due to sex-specific gene-regulation patterns associated with multiple factors (e.g. hormones, lifestyle, influences of other genes, etc), our analytic pipeline includes (with details presented in the graphical abstract):

1. Performing sex-specific association analyses (X-AS) between DNA methylation levels and phenotype across the X chromosome.
2. Testing for LVE-X at loci with statistically significant associations in females in step 1, using methods described in section 2.1.
3. Summarization of test results and visualization of DNA methylation patterns in interesting regions.

Assuming step 1 found loci of interest, we then introduce a model for detecting locus-specific variable escape from XCI (LVE-X), including whether the rate of escape depends on phenotypes and covariates (**Figure 1**). In females, we first conceptualize what may be happening in individual cells at a single CpG site, and then we aggregate the resulting probabilities across cells, since our datasets are from peripheral blood samples measuring bulk levels of DNA methylation. For a chosen CpG site and the *i*^*th*^ female participant (*i* = 1, …,*n*), each allele within cell *k* could be unmethylated (0) or methylated (1). One of the two alleles in each cell is presumed to undergo transcriptional silencing during early embryonic development as part of the XCI process; this allele is referred to as the inactive allele (*IA*), with the other allele being termed *AA*. We use this notation instead of Xi and Xa to reflect that these labels are inferred from the methylation patterns, not observed. The corresponding allelic methylation states are denoted by *Ψ*_*ikIA*_ and *Ψ*_*ikAA*_, respectively. Therefore, the methylation level for cell *k* will have potential values (0, 0.5 and 1.0) and can be written as

**Figure 1.**
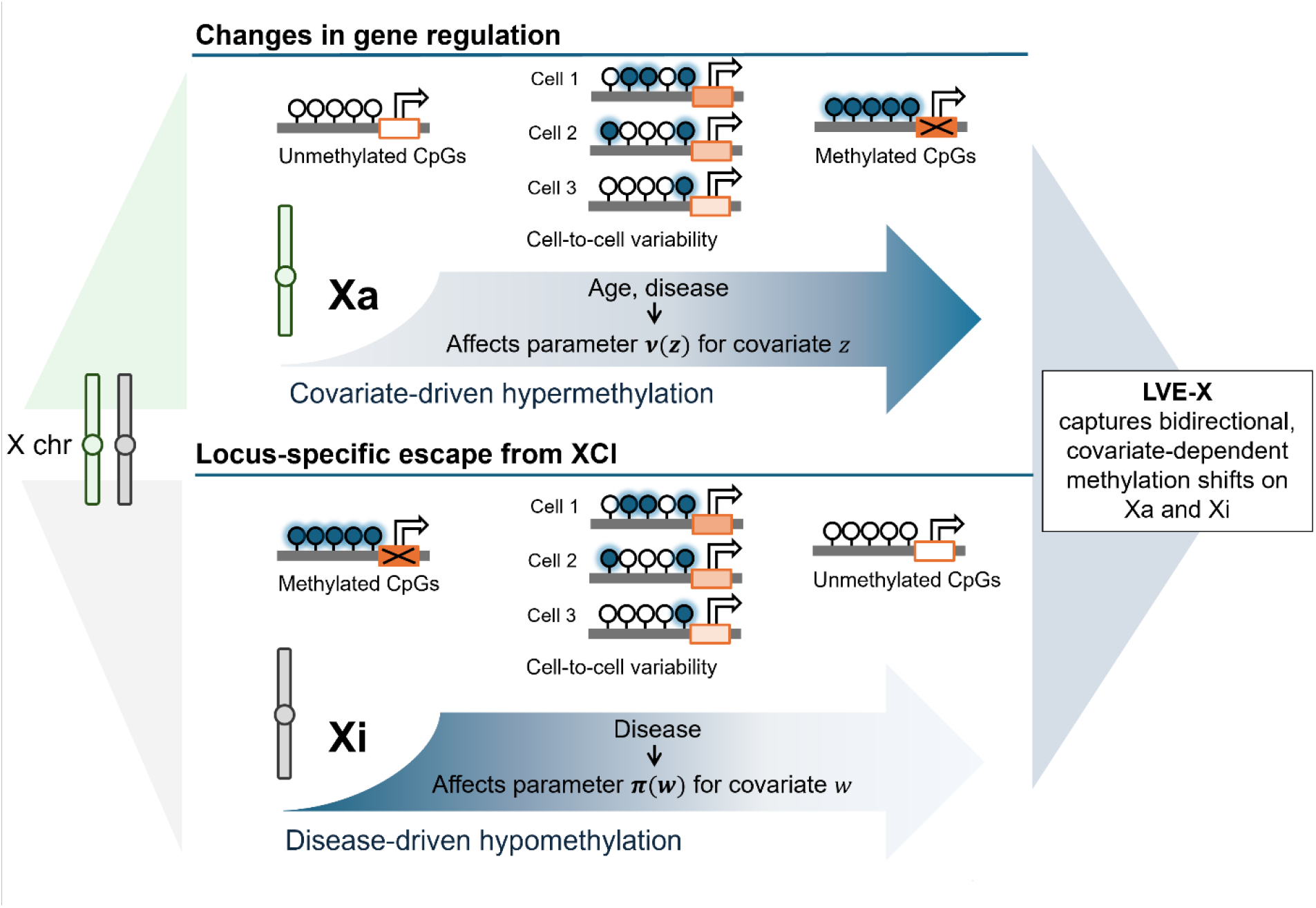
Schematic of the LVE-X model which captures locus-specific variability in X-chromosome methylation. Covariates (e.g., age, cell type composition) and phenotypic status may independently influence the methylation patterns on the active X chromosome (Xa) and inactive X chromosome (Xi). Filled and unfilled circles represent methylated and unmethylated CpGs, respectively, which allow transcriptional silencing or activity. Cell-to-cell variability is depicted for individual cells, highlighting the stochastic nature of methylation. The top panel shows covariate- or disease-driven hypermethylation on Xa (parameter *v(z)*), while the bottom panel shows disease-driven hypomethylation on Xi (parameter π(w)). These trends can be separately estimated. LVE-X integrates these bidirectional and covariate-dependent shifts to estimate locus-specific escape from X-chromosome inactivation.

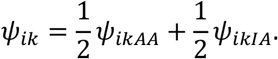

Under complete XCI, at an actively transcribed gene, the canonical expectation is that *Ψ*_*ikAA*_ = 0 and *Ψ*_*ikIA*_ = 1 . However, partial escape from XCI or alternative regulatory mechanisms such as the action of enhancers or silencers may lead to deviations from this pattern.

Therefore, we let *z*_*i*_ denote covariates that influence the probability that allele *AA* may be methylated, and *w*_*i*_ denote covariates influencing whether allele *IA* is unmethylated. To accommodate these effects, the allelic methylation states are independently modeled using Bernoulli distributions with logit link functions,

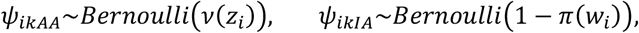

where a logistic link function specifies the means as functions of the covariates with parameter vectors *α* and *β*:

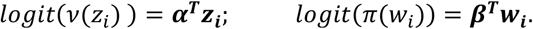

We then make an asymmetric assumption: once an allele escapes XCI it becomes subject to regulatory processes that might methylate that allele. However, we assume that the AA allele cannot be silenced through the inactivation mechanisms. Mathematically, this means *π*(*w*_*i*_) has no effect on *Ψ*_*ikAA*_. Therefore,

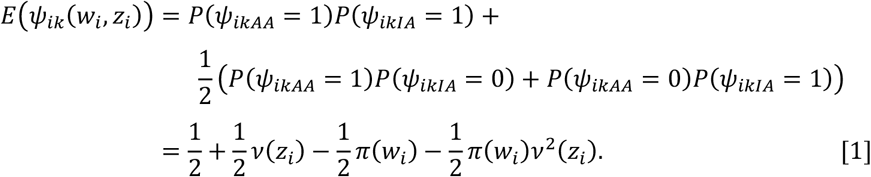

#### Bisulfite sequencing measures of DNA methylation

Given that we have data from bulk tissue samples, we assume cells are randomly selected for bisulfite sequencing. We assume that all cells from the same female share a common distribution for their methylation proportions, conditional on the covariates, *w*_*i*_ and *z*_*i*_, so that *Ψ*_*i*_(*w*_*i*_, *z*_*i*_) = *E*(*Ψ*_*ik*_(*w*_*i*_, *z*_*i*_)). For a single CpG, let *Y*^*F,D*^ denote the observed methylated read count for female *i*, where the superscript *D* indicates that the outcome is a discrete count from bisulfite sequencing data, and *F* refers to females. Let *m*_*i*_ denote the corresponding total read count. We adopt a beta-binomial (BB) model (Tripathi, Gupta, and Gurland 1994), such that

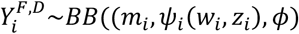

A dispersion parameter, *ϕ* ∈ (0, ∞), addresses the overdispersion in these data (Zhao et al. 2024), with smaller values corresponding to greater overdispersion relative to the Binomial distribution.

Inspection of equation 1 reveals an identifiability issue, parameter values cannot be uniquely determined if *w*_*i*_ = *z*_*i*_ with only 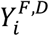 available. To address this limitation, information from male participants can be incorporated into the model. Let 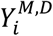 denote the observed methylated read count for male participant *i* . Since males have only one X chromosome which does not undergo XCI, only covariates *z*_*i*_ are pertinent. We assume that *v*(*z*_*i*_) is identical in females and males, conditional on the values of *z*_*i*_. We can then model 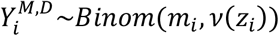 with a binomial distribution and obtain a joint log-likelihood for all participants. Additional details on the methods as well as simulations to assess performance are provided in **Supplemental Methods S1 and S2**. Analyses were conducted using a custom-developed R package called LVEXdetector which can be obtained from https://github.com/GreenwoodLab.

#### DNA methylation measured by the Illumina EPIC array

Illumina methylation array data arise from fluorescent intensity measures and are continuous rather than sequencing counts. For such data, the observed outcome at each CpG site is a methylation proportion bounded between 0 and 1. Let 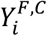for this continuous measure (*C* ) in females, and 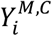 be the measure for males. For females we assume the beta distribution, 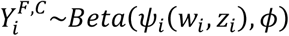, again allowing for an overdispersion parameter. Analogously to bisulfite sequencing data, male participants can contribute information to a joint modelling framework. Details are provided in **Supplemental Methods S1.3**, and the R package LVEXdetector also analyzes Illumina data.

We used two datasets to evaluate LVE-X: (i) bisulfite sequencing of stored peripheral blood samples with a custom targeted library from individuals with and without RA (the RA dataset), and (ii) DNA methylation in blood DNA with the Illumina Methylation EPIC array from rVTE patients (the rVTE dataset).

### 2.2 Description of the RA dataset

#### Data sources

The RA dataset is derived from two prior studies of stored peripheral blood samples from participants in a Quebec population biobank, CARTaGENE, that examined associations between positivity status for anti-citrullinated protein antibodies (ACPA) (Puszczewicz and Iwaszkiewicz 2011) and DNA methylation (Shao et al. 2019; Zeng et al. 2021). Given that ACPA positivity is strongly associated with RA risk, follow-up data were obtained from participants on RA diagnoses.

The CARTaGENE cohort (Awadalla et al. 2013) consists of approximately 40,000 individuals from four regions in the province of Quebec, Canada: Montreal, Quebec City, Sherbrooke, and Saguenay--Lac-Saint-Jean, recruited between 2009 and 2015. For each participant, demographic and health questionnaires were collected, and blood, serum, and urine samples were stored. Among the first 19,995 participants enrolled, stored serum samples from 3,600 individuals were analyzed for anti-citrullinated protein antibody (ACPA) levels using an INOVA enzyme-linked immunosorbent assay (Quanta Lite CCP3 IgG; Inova Diagnostics Inc., San Diego, CA, USA). Based on these measurements, two datasets were derived. In 2017, 137 individuals with either high ACPA levels (optical density > 60) or negative ACPA levels (optical density ≤ 20) were selected, and their stored blood samples underwent custom methylC-capture bisulfite sequencing (Allum et al. 2015; Shao et al. 2019). In 2019, 120 additional individuals were selected with either highly positive ACPA (optical density > 60) or negative ACPA (optical density ≤ 20), and their stored blood samples were likewise processed using targeted custom capture bisulfite sequencing, albeit with an updated targeted sequencing library (Zeng et al. 2021). Blood cell composition and genotype data for the sequenced samples were obtained from ancillary laboratory pipelines implemented during the sequencing workflow. These studies were undertaken in accordance with CARTaGENE study number 582582 and ethics board approval IRB-A04-M46-12B from the Faculty of Medicine and Health Sciences at McGill University.

#### Data description

We searched for rheumatoid arthritis followup diagnoses for any participant in these two ACPA studies. CARTaGENE asked all participants at recruitment whether they had ever received a diagnosis of RA, and then performed a follow-up questionnaire in 2016 (https://portail.cartagene.qc.ca/study/suivi_sante_2016) which asked the same question. We requested the follow-up data, and obtained approval to link the study participants to the Quebec provincial health insurance agency, RAMQ (Régie d’assurance maladie du Québec) which contains all publicly reimbursed physician services, including diagnostic International Classification of Diseases codes (ICD-10), under CARTaGENE study number 731377 and approval #2024-4041 from the CIUSSS West-Central Montreal Research Ethics Board (REB). An RA diagnosis was inferred from RAMQ code 714.x (only codes 714.0 or 714.9 were found). Report of RA from any of these three sources was considered to be “yes”, and no report of RA was considered as “no”. **Table S1** provides the source of information for each individual with RA.

From the 137 individuals selected in 2017, 15 subjects lacked cell-type composition information, and 1 subject had a low-positive ACPA-level. These 16 individuals were excluded from our analyses. Among 121 participants, 59 were ACPA positive and 62 were ACPA negative. From the 120 individuals selected in 2019, cell-type composition information was missing for 8. Therefore, 112 individuals were retained of whom 57 were ACPA positive and 55 negative (**Table 1**). RA diagnosis from RAMQ and CARTaGENE questionnaires occurred in 37 individuals (**Table 1**), of whom 24 were diagnosed after recruitment (11 in each of the two datasets; **Table S1**). As expected, ACPA+ individuals had a higher rate of RA, as expected (2017: 25.4% of ACPA+ versus 6.5% of ACPA−; 2019: 20.9% of ACPA+ versus 9.1% of ACPA−). **Table S2** presents descriptive characteristics of selected participants.

**Table 1.** Sample sizes in the RA dataset, by year of sampling from the CARTaGENE esource, ACPA status, and RA status.

|  | Total | Males | Females |
| --- | --- | --- | --- |
| **A. Number of individuals in the 2017 sampling** |  |  |  |
| Total number of participants | 121 (100%) | 43 (100%) | 78 (100%) |
| ACPA-, RA- | 58 (48%) | 21 (49%) | 37 (47%) |
| ACPA-, RA+ | 4 (3%) | 1 (2%) | 3 (4%) |
| ACPA+, RA- | 44 (36%) | 18 (42%) | 26 (33%) |
| ACPA+, RA+ | 15 (12%) | 3 (7%) | 12 (15%) |
| **B. Number of individuals in the 2019 sampling** |  |  |  |
| Total number of participants | 112 (100%) | 50 (100%) | 62 (100%) |
| ACPA-, RA- | 50 (45%) | 22 (44%) | 28 (45%) |
| ACPA-, RA+ | 5 (4%) | 3 (6%) | 2 (3%) |
| ACPA+, RA- | 43 (38%) | 19 (38%) | 24 (39%) |
| ACPA+, RA+ | 14 (12%) | 6 (12%) | 8 (13%) |
| **C. Number combining 2017 & 2019** |  |  |  |
| Total number of participants | 233 | 93 | 140 |

The bisulfite sequencing reads were re-aligned with BISMARK version 0.18.1 (Krueger and Andrews 2011), part of the genpipes v6.1.0 pipeline, to human genome build 38, and chromosome X results were retained for this study. The raw aligned data contained 1,245,869 CpG sites on the X chromosome in the 2017 data, and 1,229,858 CpG sites in the 2019 data. Quality control was performed as follows: methylated counts greater than 100 were removed from the analysis; CpG sites were then filtered again to retain only those for which at least 20 participants had a total read count greater than 2, and for which the standard deviation of the methylation proportion exceeds 0.05, resulting in 192,834 sites for analysis of the 2017 dataset and 241,127 CpG sites for the association analyses in the 2019 dataset. Of these, 175,259 sites overlapped (**Table S3**). The library design was improved between the 2017 and 2019 captures, leading to better read depth and coverage.

Proportions of 6 common blood cell subpopulations were measured in the CARTaGENE samples (neutrophils, eosinophils, basophils, lymphocytes, monocytes and granulocytes) and the first principal component of these proportions was calculated.

#### X-chromosome Association Analyses (X-AS)

We performed two primary region-based aggregation approaches and a complementary single-CpG analyses. Given the improvement in library design between the 2017 and 2019 custom captures, there were some CpGs with data in only one of the 2017 or 2019 datasets. Since gene promoter regions and CpG islands (CGI) were well captured in both libraries, our primary analyses focused (i) candidate gene promoter regions and (ii) all CpG islands on the X chromosome. Given the potential for model instability linked to low read depth (see below), as well as consequent low statistical power, we aggregated the sequencing counts in these regions. Firstly, we conducted a candidate-gene analysis, creating an aggregate measure across the promoter regions of 48 immune-related genes with potential or previous association with RA or autoimmune disease (**Table S4** lists genes and provides citations). We defined the promoter region as the transcription start site (TSS) +/- 1500bp, then summed methylated counts and unmethylated counts across all CpGs in these regions. The sums were then analyzed as a single locus measure of DNA methylation. Secondly, we analyzed all CpG islands on the X chromosomes by similarly aggregating the counts across the islands. Island definitions were obtained from the Bioconductor *annotatr* package (March 2026 download). In total, we analyzed 804 aggregated regions. For completeness, (iii) single CpG analyses were also performed in the 2019 dataset with validation of interesting loci using the 2017 dataset.

Since ACPA-positive RA may be different from ACPA-negative RA (Puszczewicz and Iwaszkiewicz 2011; Cunningham et al. 2023), sex-specific analyses were undertaken with beta-binomial regression models, at each CpG, to test for association between methylation levels and a three-group categorical variable combining ACPA status and RA status: rheumatoid arthritis–positive (RA+), ACPA-positive rheumatoid arthritis–negative (ACPA+/RA-) and ACPA-negative rheumatoid arthritis–negative (ACPA-/RA-). The latter group was used as the reference category. There were too few individuals who were RA+ and ACPA- to analyze this group separately (**Table S1**). Models included age and the first principal component of the cell type composition measures to adjust for confounding by the distribution of cell types. The decision was made to only include the first principal component, due to the small sample sizes being analyzed. In LVE-X analyses, *vv*(. ) was allowed to depend on age, the first principal component of cell type composition, and phenotypic group, whereas *π*(. ) only included phenotypic group.

Model fitting was conducted via a custom-built optimizer designed to obtain the maximum likelihood estimates of all coefficients corresponding to the covariates. Convergence was declared when either the objective function improved by less than a factor of (10^-10^ ) or the maximum number of 5,000 iterations was reached. Characteristics of the bisulfite sequencing data, such as low read depth or little inter-individual variability, led to problems with convergence at some CpGs, and **Supplemental Methods S1.4** describes the strategies taken when such problems occurred.

### 2.3 Description of the rVTE dataset

#### Data source

The rVTE dataset is derived from the REVERSE (REcurrent VEnous thromboembolism Risk Stratification Evaluation) I prospective cohort study aimed to identify patients at lower risk for VTE recurrence after completing 5–7 months of anticoagulation after a first unprovoked event (Rodger et al. 2008). All patients underwent baseline imaging with compression ultrasound for DVT and/or V/Q scan for PE to confirm the index VTE. ultrasound for DVT and/or V/Q scan for PE to confirm the index VTE. A first unprovoked VTE was defined as occurring in the absence of any of the following risk factors: leg fracture, lower-extremity plaster cast, immobilization for more than three days, surgery using general anesthetic three months prior to the index VTE, or up to five years before the time of enrollment. Individuals were excluded if they were under 18 years of age, had already discontinued anticoagulant therapy, required ongoing anticoagulation for reasons other than VTE, were geographically inaccessible for follow-up, were being treated for a recurrent unprovoked VTE or a previously known high-risk thrombophilia, defined as known deficiency of protein S, protein C or antithrombin, known persistently positive anticardiolipin antibodies (>30U mL−1), a known persistently positive lupus anticoagulant or two or more known defects (e.g. homozygous for factor V Leiden (FVL) or prothrombin gene mutation (PGM), or compound heterozygous for FVL and PGM), or were unable to consent. After stopping anticoagulant therapy, patients were followed every 6 months, and any suspected recurrences were independently adjudicated. Documented risk factors for VTE, patient-reported post-thrombotic symptoms, concomitant medications and results of thrombophilia testing were documented. Patients were censored at withdrawal, loss to follow-up, death, or re-initiation of antithrombotic therapy. Detailed description on the inclusion and exclusion criteria can be found elsewhere (Rodger et al. 2008). Recurrent VTE was defined as a new objectively confirmed VTE occurring after the index event, following completion of initial anticoagulation. Descriptive statistics of the rVTE dataset are presented in **Table S5**.

### Data description

From the original REVERSE I cohort, DNA methylation was profiled in 544 blood samples using the Illumina EPIC array (Bezerra et al. 2025), a platform that measures DNA methylation through fluorescent intensities. Quality control was performed with the R package *meffil* (Min et al. 2018), flagging samples with abnormal probe intensities, poor correlation with genotype outliers in principal component analysis, or discrepancies between reported and predicted sex. Probes with detection p-values >0.01 or fewer than 3 beads in over 20% samples were removed. Normalization followed the *meffil* pipeline, with control probes ensuring batch consistency. Post-normalization, probes overlapping SNPs with minor allele frequency >5% or known to cross-hybridize were excluded. Methylation at each CpG was quantified by the methylation proportion, ranging from 0 (unmethylated) to 1 (fully methylated). The blood cell type proportions were estimated using the EpiDISH algorithm with IDOL-optimized CpGs for reference-based deconvolution (Teschendorff and Zheng 2017). To reduce collinearity, principal component analysis was applied to the estimated proportions.

### X-chromosome-wide Association Analyses (X-AS)

After filtering, 417 participants of European descent, including 207 females (175 without recurrence, 32 with recurrence) and 210 males (141 without recurrence, 69 with recurrence), 19,517 X-chromosome CpGs were retained for analysis. Analysis of logit-transformed methylation proportions at all X-chromosomal CpG sites included on the Illumina EPIC array was conducted in females. Cox proportional hazards regression, implemented in the *survival* R package (Therneau 2026), was used to evaluate the association of each CpG with the hazard of recurrent VTE. Participants were followed from the time of first VTE until recurrence or censoring at the end of follow-up or last known contact. To prevent methylation values of −∞ or ∞ arising from logit-transformed methylation proportions of 0 or 1, extreme values were adjusted to 1×10^-6^ and (1 – 1×10^-6^). The logit-transformed methylation proportions were then computed using the *wateRmelon* R package (Pidsley et al. 2013), and used in Cox regression. The original methylation proportions were used for the LVE-X analyses. Models were adjusted for age at first VTE, the first principal component derived from the genome-wide methylation data, as well as the first principal component capturing variation in estimated cell-type proportions. Only the first principal component from both methylation and cell-type proportions was included due to model limitations on the number of covariates, and sensitivity analyses indicated comparable performance to models including additional components. No CpG reached the Bonferroni-adjusted significance threshold (p < 2.6× 10^-6^). For exploratory purposes, CpGs with p < 1×10^-4^ were considered suggestively associated, capturing the twelve strongest X-AS signals.

## 3 Results

For both datasets (RA and rVTE), we first conducted sex-specific analyses (X-AS) to identify methylation loci that were significantly associated with the pertinent phenotypes. Then, loci showing associations were further evaluated using the proposed LVE-X model. Regional visualizations of results were performed for the statistically significant loci. An overview of the pipeline is provided in the graphical abstract.

### 3.1 Analyses of methylation patterns in the RA dataset

We analyzed the summed methylated and unmethylated counts in promoter regions of 48 candidate genes and 756 CpG islands on the X chromosome. Some CpG islands were only captured in one of the two sequencing libraries; candidate gene promoter regions were always covered in both libraries. Using a Bonferroni significance threshold of 0.05/(756+48)=6.22×10^-5^, among males there were no genes and 2 CpG islands (CGIs; ChrX: 40094181-40109994 [near *BCOR*] and ChrX: 69503888-69506131 [near *NALF2*]) that showed association in either of the RA+ or ACPA+/RA- groups, compared to the ACPA-/RA-group. However, in females there were 12 genes and 144 CpG islands where at least one of the RA+ or ACPA+/RA- groups showed differential methylation from the ACPA-/RA- group. X-AS results, LVE-X and sensitivity analysis for all candidate genes and CpG islands can be found in **Tables S6.1-S6.9 and S7.1-S7.5; Figures S1 and S2** shows QQ-plots of p-values for the aggregated analyses, while **Figure S3** shows the single CpG results).

We then used the LVE-X model to estimate whether the methylation levels on the two alleles, IA or AA, were different between the three phenotypic groups in the regions selected from X-AS. Since the model decomposes the observed methylation signal into multiple latent components while also accounting for overdispersion, it is inherently more statistically demanding than a standard Binomial or Beta-Binomial model, and thus typically requires a larger sample size to achieve comparable inferential precision and power. The parameters of this model are identifiable when the locus on the male X chromosome is active, and/or when covariate influences on methylation levels differ between the inferred AA and IA alleles. **Supplemental Methods S2** contains more details on the algorithm and the results of simulation studies assessing the performance of our model for estimating LVE-X from methylation data under several scenarios. Given the assumptions, simulations show performance is good with little bias. Furthermore, simulations confirmed that incorporating information from male samples improved estimation, yielding reduced bias and lower variance in the parameter estimates.

Figure 2 illustrates our pipeline through results at four selected loci: the promoter regions of *IL13RA1, IRAK1*, and *TAF1*, and the CpG island located at ChrX:107716148-107716684 near *TSC22D3*. (Full results are in **Table S6.4 and S6.8**).Methylation levels in males are close to zero at these four loci, and in females the mean methylation levels range between about 25% and 40%. *IL13RA1* showed the strongest statistical significance in the female-specific analyses for the difference between RA+ and ACPA-/RA- groups (p=2.1x10^-27^), and **Figure 2** shows that its methylation levels are higher in RA+ females than in the other two phenotypic groups in females. *IRAK1* displayed the smallest p-value when comparing ACPA+/RA- females to ACPA-/RA- females (p=1.69x10^-19^), and the ACPA+/RA- group has the highest methylation. Twelve candidate genes showed at least one statistically significant association (p<6.22x10^-5^) in either of the two comparisons; these genes are shown in **Table2.** Among them, *TAF1* is also highlighted in **Figure 2**. In addition, one of the 144 CpG islands that passed our significance threshold was selected for display in **Figure 2**, where the ACPA+/RA- group had higher methylation than ACPA-/RA- (p=6.71x10^-8^).\

**Figure 2.**
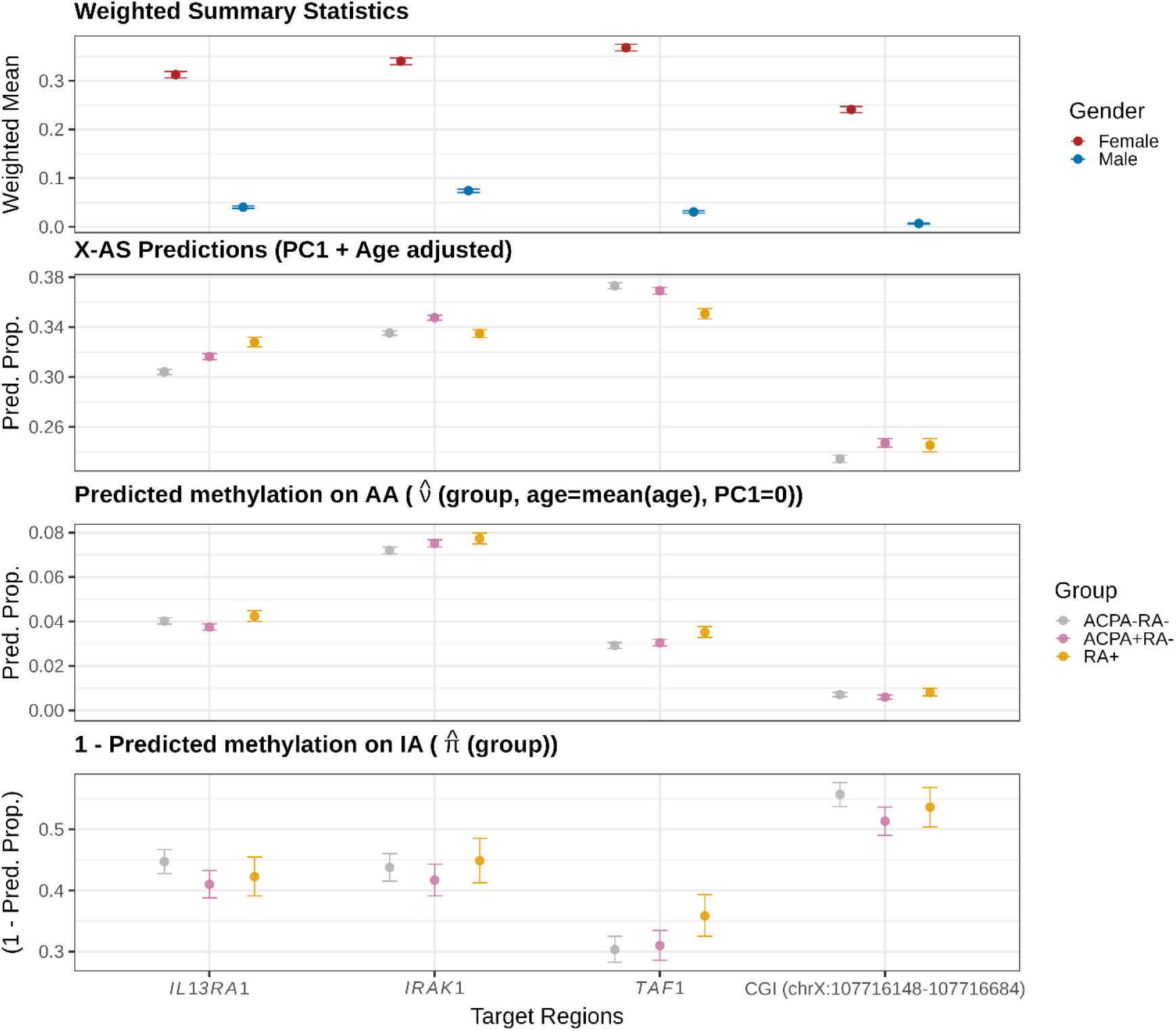
Pipeline and LVE-X model results from aggregate analyses at three candidate genes (*IL13RA1, IRAK1, TAF1*), and one CpG island (CGI; ChrX:107716148-107716684) on the X chromosome. The top panel shows mean methylation levels weighted by read depths, +/-one standard deviation, for males and females. In the second panel, results of the X-AS analysis in females are shown by the predicted mean methylation levels in three phenotypic groups defined by RA and ACPA status. For the predictions, age is set to the mean age, and the first principal component of cell type proportion is set to zero (±95% confidence intervals). In the third panel, the predicted methylation proportions (±95% confidence intervals) are shown for the AA alleles from the LVE-X model; *v*(.)is estimated when the first principal component of cell type composition is zero, and age is the mean age. In the bottom panel, one minus the predicted methylation proportions (π(.); ±95% confidence intervals) are shown for the IA alleles from the LVE-X model.

The bottom two panels of **Figure 2** show LVE-X model estimates. For the three selected genes, the estimates of *v* (. ), representing methylation levels in females on AA, are close to zero, as was seen in the males. However, at *IRAK1*, methylation levels are lower in ACPA- /RA- than in the other two groups (p=0.0024 versus RA+, p=0.004 versus ACPA+/RA-). In the bottom panel, on IA, the confidence intervals in the three phenotypic groups overlap for *IL13RA1* and *IRAK1*, indicating little evidence for differential levels of variable escape at these two genes. In contrast, at the *TAF1* promoter, containing 132 CpGs, the estimate of π(. ), one minus the estimated methylation proportion on IA, is higher in RA+ implying lower methylation levels in RA+ (**Figure 2** bottom panel, **Table S1.6**; p=1.21x10^-5^). At the CpG island selected for **Figure 2**, ACPA+/RA- individuals have lower estimates of π on IA, i.e. higher methylation levels than the ACPA-/RA- group (Table S1.8; p=0.005). Additional figures and tables for other candidate genes and CGIs are shown in **Figures S4, S5, S6** and **S7**; **Tables S6.4** and **S6.8**.

In **Table 2**, LVE-X model estimates indicate that there are six genes (*CXCR3, IL13RA1, IRAK1, RPS6KA3, TAF1, WAS*) where our model’s estimates of π are higher for RA+ than for the other groups, implying lower methylation levels on IA for RA+. There are three genes with the opposite pattern (*CNKSR2, MIR106A, XIAP*) where IA’s methylation levels are estimated to be higher in RA+. Results at *XIST* for *v*(. ) are in line with expectation for AA, with methylation levels estimated to be near 100%; despite this, the LVE-X model estimates a significant difference between RA+ and ACPA-/RA- (p=3.79x10^-4^) with higher methylation for RA+ individuals. On IA at *XIST*, all three groups have methylation levels estimated near 50-60%.

**Table 2.**
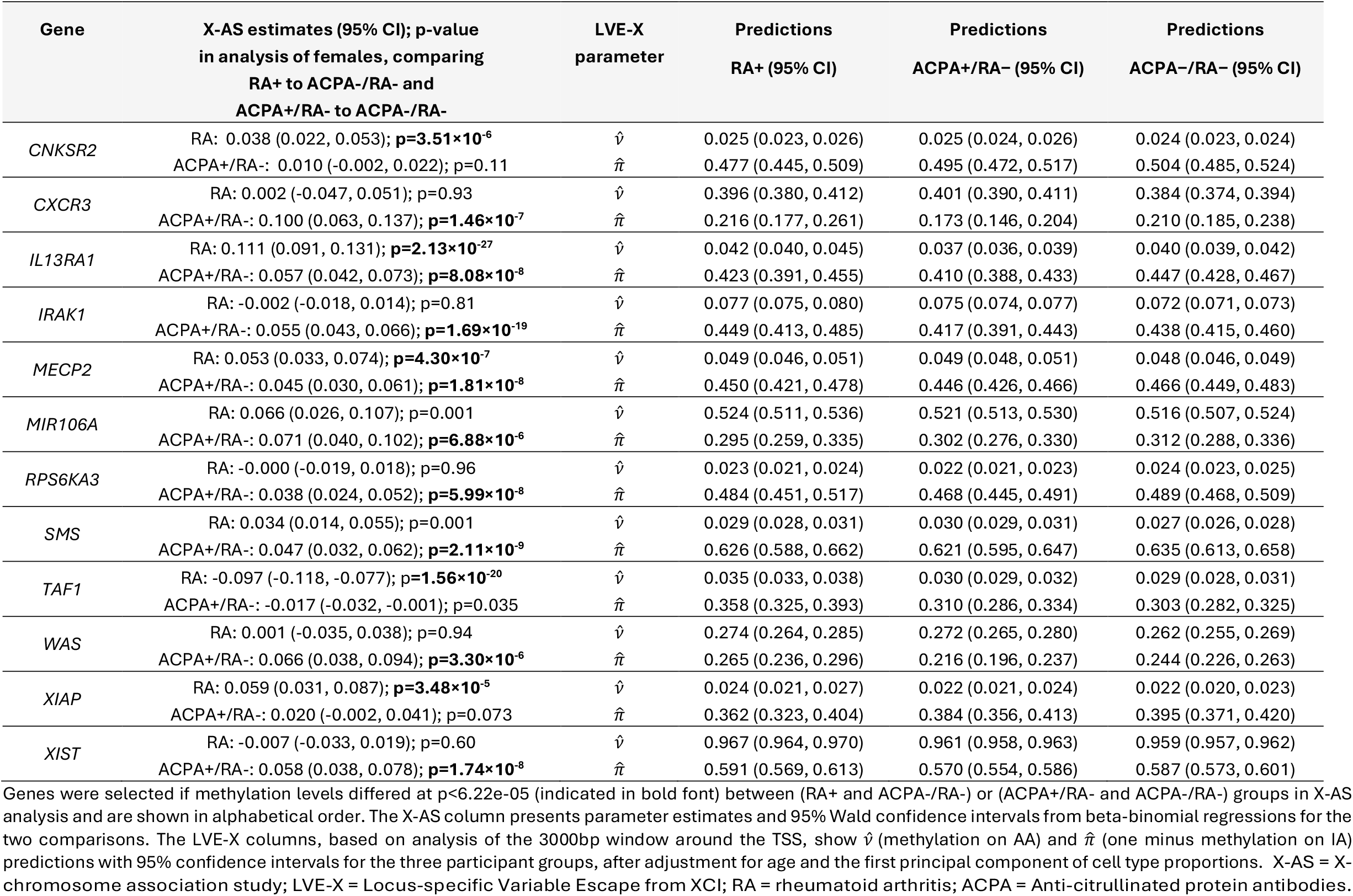
LVE-X model results for aggregated promoter methylation data in selected candidate genes associated with rheumatoid arthritis and autoimmune disease.

We also analyzed the individual CpGs for completeness, using the better-covered 2019 dataset for discovery and the 2017 dataset for replication. At many of the 241,127 CpGs in the 2019 dataset, X-AS analyses resulted in convergence issues (**Supplemental Methods S1.4**). After adjustments to address this, when possible, in females we found 22 CpG sites where methylation levels differed at p<0.0001 between RA+ and ACPA-/RA-, and 25 CpG sites when comparing ACPA+/RA- to ACPA-/RA-. Among males, 7 CpG sites were identified in each of the two comparisons at p<0.0001; none of these identified 61 CpGs overlapped. Although a Bonferroni-corrected threshold for 241,127 tests would be 2.07x10^-7^, or smaller if we corrected for four comparisons, we used a less stringent p-value threshold given that sample size and read depth limit our statistical power. In the 2017 dataset, focused validation identified only 1 of these 61 CpGs with p<0.05. The single CpG results with p<0.0001 can be found in **Tables S6.10-S6.15** and **S7.6-S7.11** and QQ-plots are in **Figure S3**. We calculated post-hoc power using the estimates from the 2019 data with the read counts from the 2017 data, and it was often poor (see **Supplemental Methods S3**, for post hoc power calculations and results). Although few CpGs showed strongly significant findings, often there were several nearby CpGs showing association at p<0.05.

The time of diagnosis was unknown for some of the CARTaGENE participants with self-reported RA. Therefore, we conducted a sensitivity analysis in which individuals with self-reported RA at the time of enrollment into the CARTaGENE study were excluded from analyses. In these analyses of incident RA diagnoses only, we refit models for the loci previously identified in the RA dataset X-AS. The full results of these sensitivity analyses for the combined data are provided in **Tables S6.3** and **S7.3;** most significant associations were retained with two exceptions: the X-AS analysis of *SMS* no longer passed our significance threshold, and at *XIST*, the comparison between RA+ and ACPA-/RA- for AA was no longer significant.

### 3.2 Analyses of DNA methylation patterns and their associations with recurrent VTE

We repeated a similar analytic strategy to illustrate X-chromosome methylation patterns and their potential associations with rVTE, illustrating that the new methods can be used for Illumina EPIC measures of DNA methylation. Consistent with previous reports, recurrence was more frequent in males than in females (32 % vs 15 %) **(Table S5)**. Males were slightly older at their first VTE and had a shorter median time to recurrence compared with females. For a complete description of this dataset, see Bezerra et al. 2025. In REVERSE I, 19,517 X-linked probes passed quality control. Among 455 unprovoked VTE participants recruited at first VTE, survival analysis for time to recurrence was fit for 207 females, adjusting for age, first genome-wide methylation principal component, and first cell-type principal component. As with the RA dataset, a p-value threshold was used to identify suggestive associations due to limited statistical power due to the rarity of recurrence in females. QQ plots and a regional X-AS plot are shown in **Figure S8**.

The CpG sites most strongly or suggestively associated with rVTE are summarized in **Table 3**. Notably, two loci mapping to *BCOR* and *UBA1* have previously been implicated in thromboembolic outcomes in genetic association studies (Poulter et al. 2021). For the LVE-X analysis, we first assessed sex differences in methylation and prioritized CpGs showing at least a 10% difference between males and females, with intermediate methylation levels in females and hypomethylation in males. We also examined CpGs within 5 kb of the associated site from **Table 3**, prioritizing those in regulatory or promoter regions, and selected *GAB3, GRIA3*, and *UBA1* for further presentation. Although most CpG correlations occur within 1 kb, they may extend up to 10 kb (Liu et al. 2014). X-chromosome regional plots show how males and females differ in methylation patterns across each CpG for these selected genes (**Figure S9)**. For the top CpG cg10400716, in the body of *GAB3*, the signal lies within a transcription factor binding peak, likely corresponding to a CTCF binding site; however, methylation proportions were very low in both sexes. We therefore focused on the TSS1500 cg02225164, 2.5kb from the associated site, showing 12% methylation in females and essentially zero in males (**Figure S9)**. Estimates of methylation on *Xa*,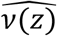, are slightly lower in rVTE, whereas the probability of escape,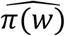, is slightly higher in rVTE, compared to those with only a first VTE (**Table 3; Figure 3**). The same patterns were observed for the CpGs at genes *GRIA3* and *UBA1*. As these differences are minimal, LVE-X likely does not differ between rVTE and VTE, suggesting that the observed association with rVTE is independent of XCI escape.

**Table 3.**
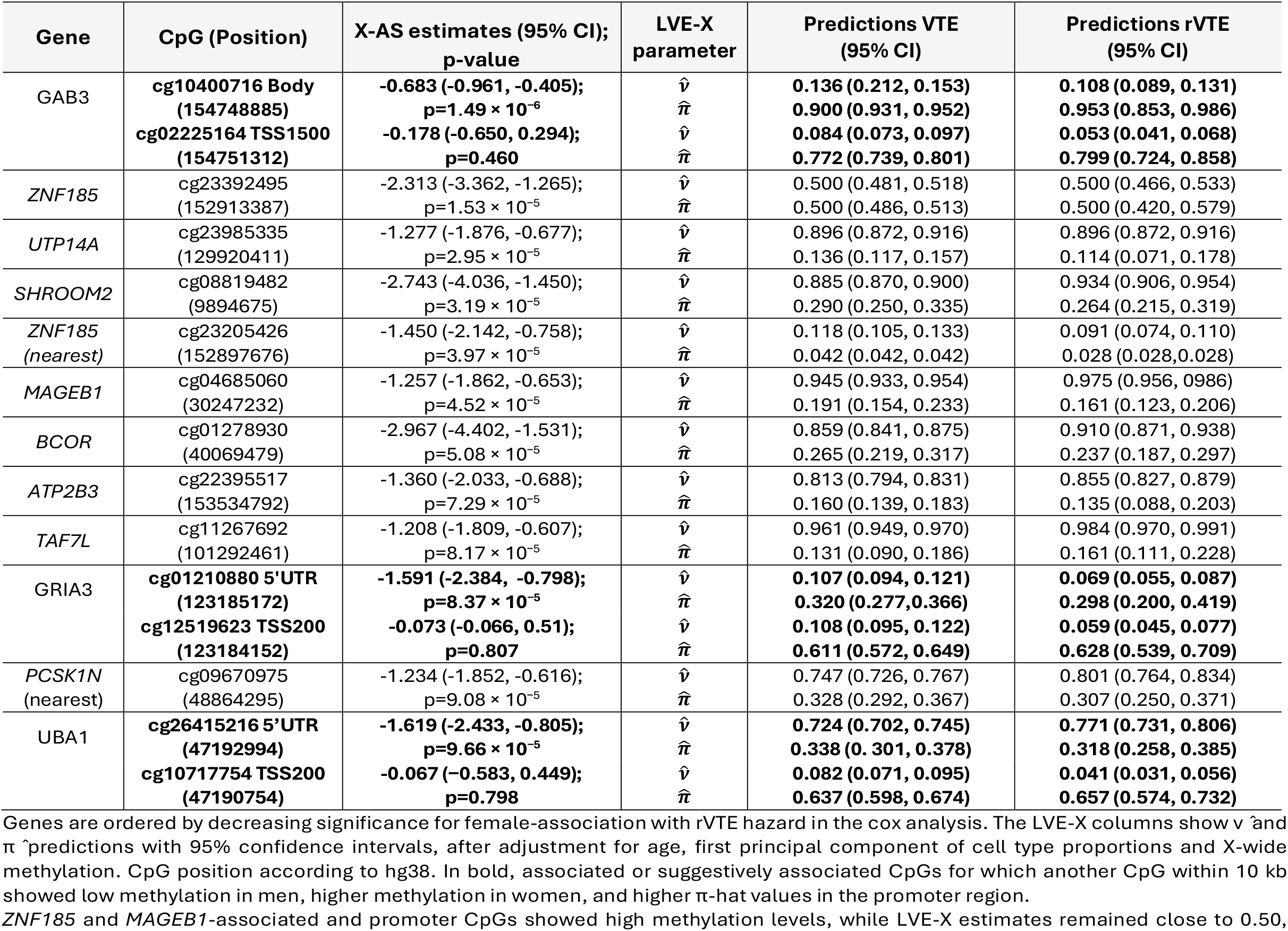
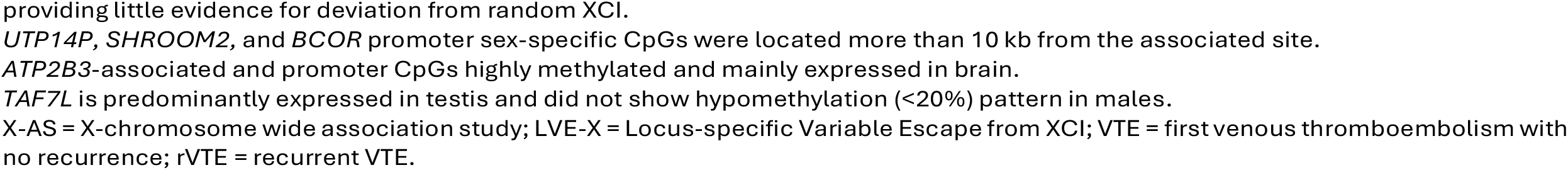
LVE-X model in CpG sites associated with recurrent VTE.

**Figure 3.**
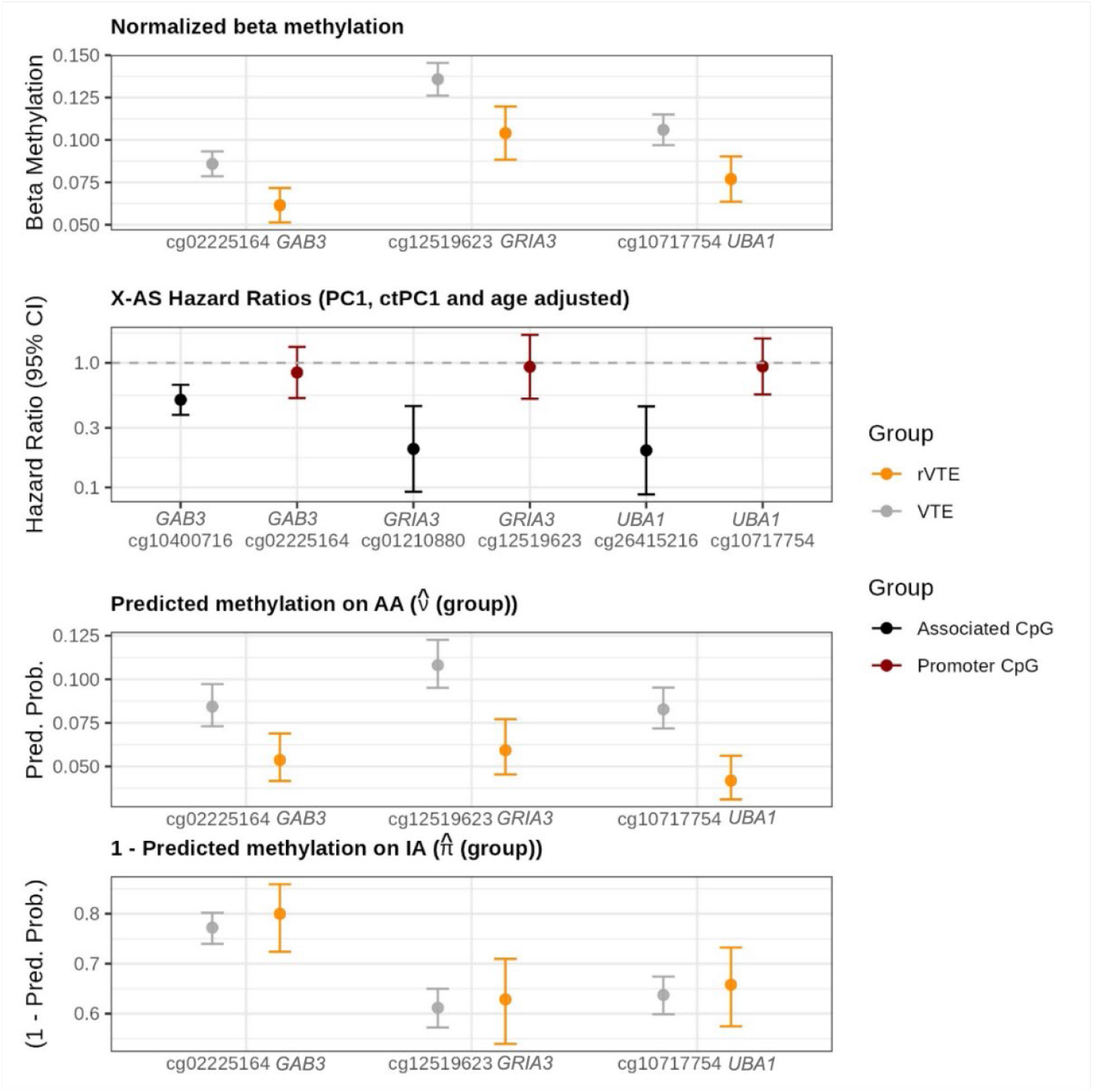
Pipeline and LVE-X model results in the rVTE dataset. CpG cg02225164 is located 2.5 kb from the top-associated CpG in *GAB3* within the TSS1500 region (-200 to -1500 bp upstream of the transcription start site). In *GRIA3*, cg12519623 (TSS200) is 1.0 kb from the associated cg01210880 (5′UTR and in *UBA1*, cg10717754 (TSS1500) is 2.2 kb from cg26415216. The top panel shows the mean methylation proportions in males and females with (±95% confidence intervals). The second panel presents hazard ratios for a one unit change in methylation associated with rVTE from female-only Cox survival analysis models, with genome-wide methylation PC1 and cell-type composition PC1 set to zero, and age fixed at the mean (±95% confidence intervals). In the third panel, the predicted methylation proportions (±95% confidence intervals) are shown for the AA alleles from the LVE-X model; *v*(.)is estimated when the first principal component of cell type composition is zero, and age is the mean age. In the bottom panel, one minus the predicted methylation proportions (π(.); ±95% confidence intervals) are shown for the IA alleles from the LVE-X model.

## 4 Discussion

The X chromosome has often been ignored or forgotten in genetic and genomic analyses despite its important role in sex-biased diseases. Although ample evidence suggests that gene escape from XCI may be associated with sex-biases, there was no statistical method to test for association between LVE-X and disease phenotypes or covariates. Since DNA methylation is a key mechanism maintaining gene silencing, and given that DNA methylation can be easily measured on large numbers of individuals, we developed a test (and a software package, LVEXdetector) for covariate-associated changes in DNA methylation patterns consistent with LVE-X. We assessed performance of this method through simulation studies and analysis of two sex-biased traits: RA and rVTE. In addition to identifying new CpGs associated with RA and rVTE through sex-specific analyses of the X chromosome data, we found loci where LVE-X is one plausible explanation for the DNA methylation patterns and sex-biased disease risk. At other loci, we ruled out LVE-X as a potential explanation, suggesting other mechanisms are driving the differential methylation.

LVE-X has a more general statistical capability. It estimates a distribution of DNA methylation levels where one allele in females matches that in the males, and the other allele may demonstrate a different distribution. We also identified loci where locus-specific escape from X-inactivation is unlikely, but where phenotype associations in males and in one of the two female alleles appear to differ from the other female allele, suggesting an association with a gene subject to undisturbed XCI. At several of the CpG islands listed in **Supplement Table S6.6**, we infer the existence of different covariate relationships between the AA and IA alleles.

Several loci were identified in our analysis of the RA data. Higher DNA methylation was found in the *IL13RA1* promoter (ENST00000371666.8) in RA+ females, which may imply lower expression of this gene. Since our analyses adjusted for cell type composition, it is unlikely that the association is driven by differences in cellular proportions. *IL13RA1* encodes the alpha-1 subunit of the Interleukin-13 receptor, which mediates signaling by IL-13, a key cytokine involved in immune regulation, inflammation, and T-helper cell polarization that is elevated in RA synovial fluid. IL-13 downregulates inflammatory processes in RA and may slow disease progression(Iwaszko, Bialy, and Bogunia-Kubik 2021; Yang et al. 2025, 2020). In this context, reduced *IL13RA1* expression, as suggested by the higher promoter methylation in RA+ females, could impair IL-13–mediated anti-inflammatory signaling and contribute to persistent inflammation. In addition, *IL13RA1* variably escapes XCI (Balaton and Brown 2021; Marcelis et al. 2025), supporting the plausibility of sex-specific epigenetic dysregulation at this locus. Although our LVE-X analyses showed little evidence for differential variable escape at *IL13RA1*, altered promoter methylation, rather than altered escape dynamics, may be a mechanism linking *IL13RA1* to RA pathophysiology. Higher methylation at the *IRAK1* promoter was also observed in ACPA+/RA− females. This is notable because ACPA positivity often precedes clinical RA and may reflect an early phase of autoimmune dysregulation. *IRAK1* is a key mediator of Toll-like receptor and Interleukin-1 receptor signaling, with established roles in promoting pro-inflammatory cytokine production. *IRAK1*-dependent signaling in synovial fibroblasts amplifies neutrophil and monocyte joint recruitment, sustaining local inflammation (Hoyler et al. 2022). In this context, epigenetic regulation of *IRAK1* may modulate the magnitude or timing of innate immune activation in early RA.

We inferred partial escape at *TAF1* downstream of the TSS in RA+ females. *TAF1* encodes a core component of the transcription factor IID (TFIID) complex and plays an essential role in RNA polymerase II transcription initiation. In multiple sclerosis, *TAF1* dysfunction can increase expression of pro-inflammatory genes and contribute to autoimmunity (Rodriguez-Lopez 2025). Furthermore, loss of *TAF1* alters differentiation-associated genes implicated in autoimmune thyroid disease and lupus erythematosus, both female-biased autoimmune conditions (Liu et al. 2025). By influencing hematopoietic differentiation and transcriptional regulation, *TAF1* dosage may broadly impact immune tolerance mechanisms relevant to autoimmunity.

The X-linked associations with rVTE highlighted genes involved in immune signaling, vascular biology, and protein regulation, with plausible roles in VTE pathogenesis. *GAB3*, the only gene reaching Bonferroni significance in females, exhibited hypomethylation in rVTE. *GAB3* encodes an adaptor protein essential for cytokine- and growth factor–mediated signaling and plays a key role in immune regulation and macrophage differentiation (Wolf et al. 2002). DNA methylation changes or escape at this gene could potentially influence VTE risk by promoting inflammation-induced coagulation and activation of endothelium. Among the identified loci, *UBA1* emerges as particularly compelling due to its previous association with VEXAS-related VTE and evidence suggesting escape from XCI. Genetic variation at this locus may influence ubiquitin-mediated inflammatory pathways implicated in thrombotic disease, consistent with the role of pathogenic *UBA1* mutations in VEXAS syndrome, a condition characterized by systemic inflammation, hematologic abnormalities, and elevated VTE risk (Poulter et al. 2021). Moreover, *UBA1* is reported to escape XCI (Balaton and Brown 2016), and our method indicated a high probability of variable escape at this locus, although this might not directly impact rVTE. Instead, the observed statistical associations, at least for rVTE, are likely to reflect alternative regulatory mechanisms, such as methylation on the active X chromosome or cell-type specific chromatin accessibility.

The LVE-X model has several important limitations when used to analyze DNA methylation levels measured in whole blood. Firstly, estimates derived from blood will miss associations in other tissues that may be more relevant to RA or rVTE. Secondly, LVE is known to be cell-type specific and may vary among immune cell types within the same individual, contributing unequally to the overall escape dosage within a tissue. For example, a study on two identical twins showed that *GAB3* exhibits escape specifically in T-CD4^+^ cells (Zito et al. 2023). In systemic lupus erythematosus (SLE), a disease with nine-fold higher prevalence in females, T-cells (Syrett et al. 2019), dendritic cells (Barrat and Guery 2025) and B-cells (Sierra et al. 2025) have changes in *Xi* activation/inactivation under certain conditions. Since our two datasets measured methylation from whole blood samples containing a mixture of cell types, estimated LVE proportions are averaged across cells, potentially obscuring cell-specific or cell-type-specific differences. Thirdly, the cell type composition in the blood samples may confound relationships we identified. All analyses were adjusted for cell type composition by using at least one principal component of the measured or estimated cell type proportions. Sensitivity analyses in the rVTE data led to similar results with different numbers of principal components. Nevertheless, further validations would be important to pursue.

One strength of the LVE-X method is that it can analyze DNA methylation data from either bisulfite sequencing or Illumina arrays: the algorithm for sequencing-derived measures is built on a beta-binomial model, whereas for Illumina data we used the (continuous) Beta distribution. For the latter, convergence was generally stable. However, convergence was not always achieved for the bisulfite sequencing data, and the issue was exacerbated by low read depths. In such cases, we fit regular binomial models, and also explored fitting some zero-inflated beta binomial models **(Supplemental Methods S1.4)**, but the problem often persisted. As a result, we analyzed aggregated counts across promoter regions and CpG islands; convergence problems were rare in these analyses. Hence, some smoothing or aggregation of data across CpGs is likely required for LVE-X to find signals of interest in bisulfite sequencing data, whereas Illumina data do not require such strategies. On the other hand, bisulfite sequencing can provide better coverage of the methylation levels along the chromosome and can therefore better localize the regions where altered methylation patterns are occurring.

The methods developed here apply to DNA methylation data from bulk tissues such as blood samples. However, the theory builds on expectations for single cells. It would be possible to extend these statistical concepts to build a single cell model for estimating locus-specific escape from XCI in DNA methylation data, which would make it possible to estimate cell-type-specific LVE-X. Furthermore, there may be potential to improve the sensitivity of our LVE-X analysis by implementing regional smoothing, as we have previously done for autosomal data (Zhao 2022; Zhao et al. 2024).

## Graphical abstract

Workflow of the proposed analyses. After quality control steps, we performed sex-specific association analysis for all CpGs on the X-chromosome, followed by our new method for assessing locus specific escape from XCI (LVE-X).

## Supplemental Methods

### S1 Supplemental information for the LVE-X model

#### S1.1 Proof of 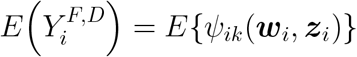

In the main text, we define 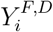 as the observed methylated read count for female participant *i* at a given CpG site, where the superscript *D* indicates that 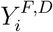 is a discrete count obtained from bisulfite sequencing data.

Since bulk tissue sequencing entails cellular lysis, intact cells are destroyed and the resulting reads originate from individual DNA molecules (alleles) rather than from whole cells. Assuming that, across cells, DNA molecules from each allele are randomly sampled for sequencing, we show that the expected observed methylation proportion for female participant

*i* at a given CpG site, 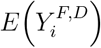, that is, the allele-averaged information for participant *i* at this CpG site, equals the corresponding model-based expectation *E*{*ψ*_*ik*_(***w***_*i*_, ***z***_*i*_)}.

To show 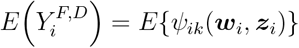, by the law of total expectation,

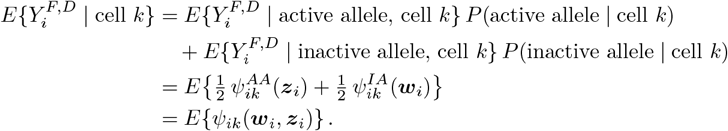

Moreover,

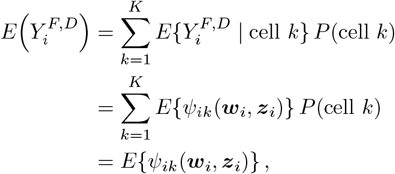

because 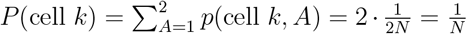, where *P* (cell *k*) denotes the probability of a certain cell *k* is selected, *N* denotes the number of cells and *A* denotes the allele-status variable. It follows that 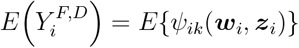, as required.

#### S1.2 Explicit Log-Likelihood for the XCI Status Detection Model for Bisulfite Sequencing Data

An explicit expression for the log-likelihood function is given by

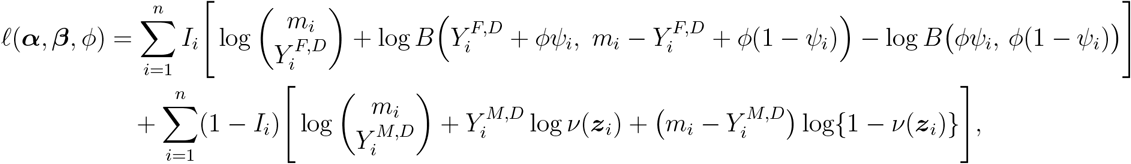

where *I*_*i*_ = 1 for a female individual, *ψ*_*i*_ = *ψ*_*i*_(***w***_*i*_, ***z***_*i*_) and *B*(·, ·) denotes the Beta function.

#### S1.3 Explicit Log-Likelihood for the XCI Status Detection Model for Illumina data

In the main text, we define 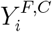 and 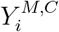 as the observed methylation level for female and male participant *i*, respectively, at a given CpG site. The superscript *C* indicates that the observed methylation level is a continuous variable from Illumina platform.

An explicit expression for the joint log-likelihood is given by

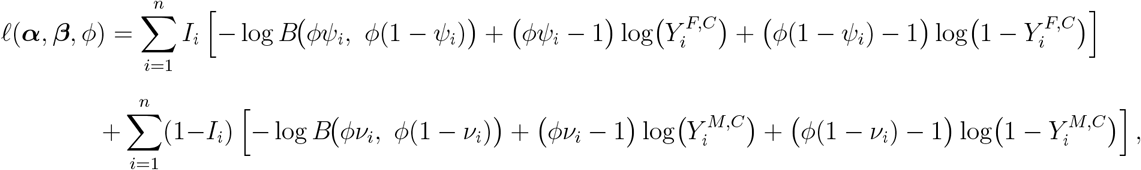

where *I*_*i*_ = 1 for a female individual, *B*(·, ·) denotes the Beta function, and *ψ*_*i*_ = *ψ*_*i*_(***w***_*i*_, ***z***_*i*_) is defined as in Equation (1).

#### S1.4 Convergence problems in the analysis of RA datasets

At some CpGs, convergence was not obtained when performing the X-chromosome association analysis separately by sex, using beta-binomial regressions. Steps taken are listed here.

1. *The beta-binomial dispersion parameter tended towards infinity, indicating no evidence for overdispersion*: In such cases, regular binomial regression models were fit instead. In the X-AS analyses of the 2019 dataset, binomial regression was applied to 103,822 sites in the female-only RA analysis, and to 147,478, 102,353, and 147,480 sites in the male-only RA, female-only ACPA, and male-only ACPA analyses, respectively. In the X-AS analyses of the 2017 datasets, we fitted binomial regression models at 70,847 CpG sites in the female-only RA analysis, 119,382 sites in the male-only RA analysis, 70,707 sites in the female-only ACPA analysis, and 111,753 sites in the male-only ACPA analysis.
2. *Lack of convergence due to separation or quasi-separation of the data by the phenotypic covariate (either ACPA status or RA status) leading to the appearance of extremely small p-values*: we examined boxplots of the methylation proportions by phenotype, and identified CpGs where there were a small number of individuals whose methylation proportions differed from the majority. Such CpGs were removed from the set of CpGs considered interesting for follow-up analyses. Specifically, there are 22,356 and 5,893 such sites in the female-only and male-only RA analyses, respectively, and 26,028 and 12,251 such sites in the female-only and male-only ACPA analyses of the 2019 dataset, respectively. In the 2017 dataset, we identified 5,594 such sites in the female-only RA analysis, 3,316 in the male-only RA analysis, 6,946 in the female-only ACPA analysis, and 11,111 in the male-only ACPA analysis.
3. *Over representation of full or zero methylation in samples:* For sites where at least 10% of the samples are either fully methylated or fully unmethylated, we considered whether a zero-inflated model could better model the actual methylation patterns in the sex-specific analyses in the situations where the previous beta-binomial model did not converge. Therefore, in the 2019 dataset, we fit zero-inflated beta-binomial models to 14,555 such sites in the female-only RA analysis, 1,957 sites in the male-only RA analysis, 16,359 sites in the female-only ACPA analysis, and 1,482 sites in the male-only ACPA analysis. With this approach, we fit a zero-inflated model when there were more individuals with zero methylation values than individuals with methylation values near 100%. When the converse was true, we fit zero-inflated models to one minus the methylation proportions. As for the beta binomial model fits, we reduced from the zero-inflated beta-binomial to the zero-inflated binomial model when dispersion estimates were greater than 1000. A zero-inflated model passes with no convergence issues for 61 sites in the female-only RA analysis, 409 sites in the male-only RA analysis, 261 sites in the female-only ACPA analysis, and 731 sites in the male-only ACPA analysis. However, no additional associations with RA or ACPA were identified at *p <* 0.0001.

#### S1.5 Parametric bootstrap for stable variance estimation

In some loci, the observed methylation proportions can be at or near the boundary (0 or 1), which may lead to a nearly singular observed Fisher information and, consequently, unstable numerical inversion when estimating the covariance matrix of the LVE-X parameter estimates. In these settings, standard error estimates obtained from the inverse information matrix can be unreliable. To improve numerical stability, we use a parametric bootstrap to estimate the covariance matrix of the estimators (Van Der Laan & Bryan, 2001). By default, our R package draws 300 bootstrap samples.

#### S1.6 Testing whether LVE-X can be excluded as an explanation

In this section, we develop a likelihood ratio test (LRT) to assess whether the phenotype is associated with locus-specific variable escape from X-chromosome inactivation (LVE-X). Specifically, letting *β*_pheno_ denote the regression coefficient for the phenotype in the model for the LVE-X probability, we test

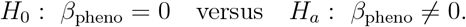

Under *H*_0_, the LVE-X probability does not depend on phenotype (conditional on any other covariates included in the model), whereas under *H*_*a*_ it varies with phenotype.

More generally, one may consider the global null hypothesis

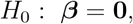

which corresponds to a model in which the LVE-X probability is constant across individuals (i.e., no covariate effects), thereby representing the absence of covariate-dependent LVE-X.

We now derive the test statistic for the hypothesis concerning the phenotype effect *β*_pheno_. The same construction extends directly to joint tests of ***β*** by imposing the corresponding set of linear constraints.

Let *l*(***θ***) denote the (joint) log-likelihood of the LVE-X model, where

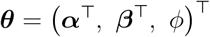

collects the regression parameters for *v*_*i*_ = *v*(*z*_*i*_) (Xa), the regression parameters for *π*_*i*_ = *π*(*w*_*i*_) (Xi), and the dispersion parameter *ϕ >* 0. We fit the unrestricted model under *H*_*a*_ to obtain the maximum log-likelihood

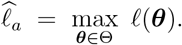

To fit the restricted model under *H*_0_, we impose the linear constraint *β*_pheno_ = 0 (equivalently, remove the phenotype term from *w*_*i*_) and maximize the log-likelihood over the restricted parameter space:

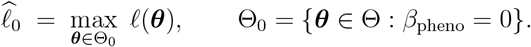

Importantly, although our likelihood may be constructed using both female and male samples, the male contribution depends only on *v*(*z*_*i*_) and not on *π*(*w*_*i*_). Therefore, the LRT comparing *H*_0_ and *H*_*a*_ is driven by the female data, while still allowing stable estimation of shared parameters through joint fitting.

The LRT statistic is

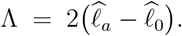

Under standard regularity conditions, 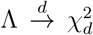 under *H*_0_, where d is the number of constrained regression coefficients (here *d* = 1 for a single phenotype coefficient; for a multilevel phenotype, *d* equals the number of corresponding indicator coefficients). The resulting *p*-value is

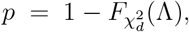

where 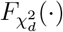 denotes the CDF of a 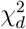 distribution. A small *p*-value provides evidence that the phenotype is associated with LVE-X (i.e., *π*(*w*) varies with phenotype). Conversely, failure to reject *H*_0_ indicates that phenotype-dependent LVE-X is not supported, and thus LVE-X is unlikely to be a primary explanation for the observed phenotype–methylation association at the locus.

### S2 Results from simulation studies to evaluate the LVE-X model performance

In this section, we evaluate the performance of the proposed models through simulation studies. We begin by assessing the model specified in Equation (2) of the Methods section, defined as

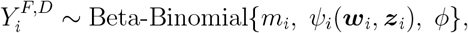

Where 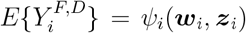, and *ϕ* ∈ (0, ∞) denotes the overdispersion parameter, with smaller values indicating greater variability relative to the standard Binomial model.

This model is fitted using data from female participants only. To ensure identifiability in this setting, the covariate vectors ***w***_*i*_ and ***z***_*i*_ are specified to differ.

In the second simulation scenario, we evaluate the model given in Equation (3), which jointly incorporates data from both male and female individuals. In this setting, it becomes feasible to use the same covariates for ***w***_*i*_ and ***z***_*i*_. The corresponding log-likelihood function is given by

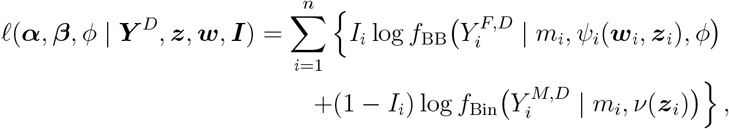

where *I*_*i*_ = 1 if individual *i* is female and *I*_*i*_ = 0 if male. Here, *f*_BB_(·) and *f*_Bin_(·) denote the Beta–Binomial and Binomial probability mass functions, respectively.

#### S2.1 Simulation for Bisulfite Sequencing Data Model

##### S2.1.1 Simulation Scenario 1

To mimic the bisulfite sequencing data-generating mechanism, we adopt the following simulation design. At a given CpG site, each cell is assumed to fall into one of three possible methylation states: no methylated allele, one methylated allele, or two methylated alleles. Accordingly, the cell-level outcome is generated using a Dirichlet–Multinomial distribution with three categories corresponding to these three potential methylation states.

The class probabilities are specified according to Equation (1) from Methods Section in the main text. The dispersion parameter in the Dirichlet–Multinomial distribution is set to 10 to induce a moderate degree of overdispersion. The total number of cells is generated from a Poisson distribution with mean 1000. For each simulated dataset, before downsampling, the methylation counts are first obtained at the cell level and then summed over cells across the three methylation states to yield the total methylated and unmethylated counts.

To mimic the downsampling inherent in sequencing experiments, an additional subsampling step is introduced. Specifically, a Hypergeometric distribution is used, where the population size is taken to be the total number of alleles across all cells (equal to twice the number of cells), and the number of “successes” corresponds to the total number of methylated alleles obtained from the Dirichlet–Multinomial component. A sample of 50 reads is then drawn from this Hypergeometric distribution, yielding the observed total read count and methylated read count used in the simulated datasets. An illustrative figure depicting the data-generating process is provided in Figure S2.1.

The simulation study is replicated 200 times under three distinct configurations of the true coefficient values,

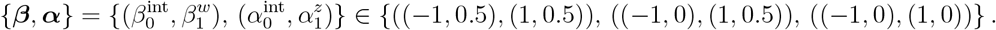

Here, 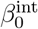 and 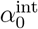 denote the intercept terms, while 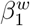 and 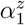 represent the coefficients associated with the scalar covariates *W* and *Z*, respectively. The covariates *W* and *Z* are independently generated from standard normal distributions. The sample size is fixed at 200. Across these replicates, model performance is summarized using the following metrics: the average bias (avgBias), the average empirical standard error (avgSEE), the average model-based standard error (avgSEM), and the average coverage rate (avgCR) of the 95% confidence intervals for the estimated model parameters.

It can be seen from Table S2.1, which summarizes the simulation results for this scenario, that the model performs reasonably well according to the reported criteria. However, the standard errors for the estimates of ***α***, in both avgSEE and avgSEM, are larger than those for the estimates of ***β***.

**Figure S2.1.**
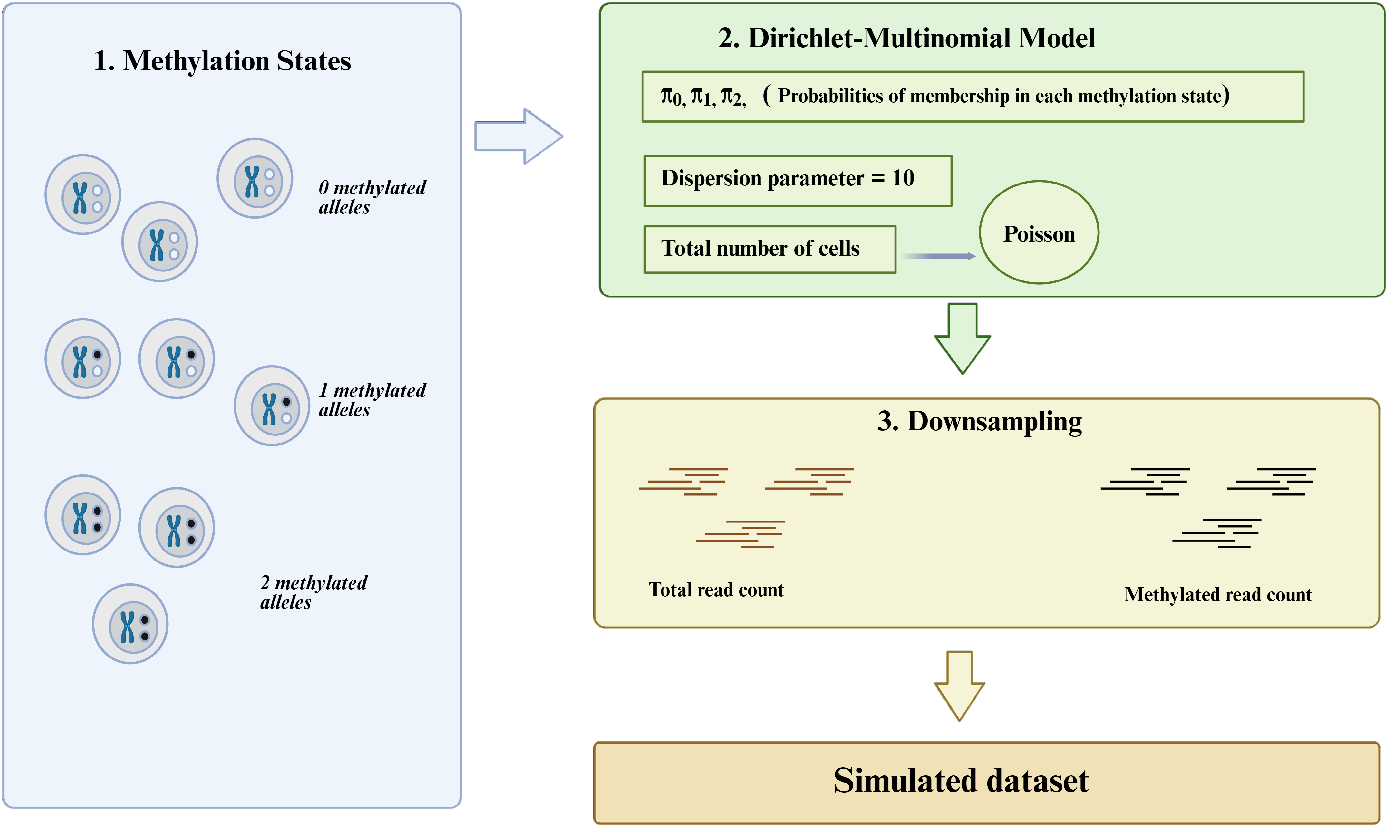
Workflow of the simulation data-generating process for bisulfite sequencing with Hypergeometric downsampling.

**Table S2.1.** Finite-sample performance in Simulation Scenario 1 (female-only bisul-fite sequencing–based model) under three configurations of the true parameter values 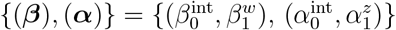.

| Truth $\{(\beta), (\alpha)\}$ | Metric | $\hat{\alpha}_0$ | $\hat{\alpha}_1$ | $\hat{\beta}_0$ | $\hat{\beta}_1$ |
| --- | --- | --- | --- | --- | --- |
| $\{(-1, 0.5), (1, 1)\}$ | avgBias | -0.097 | 0.039 | -0.122 | 0.055 |
|  | avgSEE | 0.569 | 0.329 | 0.266 | 0.142 |
|  | avgSEM | 0.506 | 0.297 | 0.229 | 0.124 |
|  | avgCR (%) | 87.5 | 91.0 | 92.0 | 94.5 |
| $\{(-1, 1), (1, 1)\}$ | avgBias | 0.022 | 0.053 | -0.030 | 0.030 |
|  | avgSEE | 0.405 | 0.256 | 0.191 | 0.154 |
|  | avgSEM | 0.386 | 0.262 | 0.163 | 0.138 |
|  | avgCR (%) | 91.0 | 94.5 | 90.0 | 92.5 |
| $\{(-1, 1), (1, 0.5)\}$ | avgBias | 0.079 | 0.057 | -0.004 | 0.011 |
|  | avgSEE | 0.576 | 0.308 | 0.195 | 0.166 |
|  | avgSEM | 0.488 | 0.234 | 0.206 | 0.165 |
|  | avgCR (%) | 93.0 | 96.0 | 90.0 | 92.0 |

##### S2.1.2 Simulation Scenario 2

In this simulation, male data are incorporated in addition to female data. Female observations are generated as in Section S2.1.1, under the same three sets of true coefficient values, but with the female sample size set to 100 and the male sample size set to 100. Male methylated read counts are then generated from a Binomial model with mean given by *v*(***z***_*i*_), using the same true coefficient vector ***α*** as for females to satisfy Assumption 1, and with the total read count fixed at 50. *Z* is also generated from a standard normal distribution. Model performance is summarized using the average bias (avgBias), the average empirical standard error (avgSEE), the average model-based standard error (avgSEM), and the average coverage rate (avgCR) of the 95% confidence intervals for the estimated model parameters.

**Table S2.2.** Finite-sample performance in Simulation Scenario 2 (Illumina sequencing–based model incorporating both female and male participants) under three configurations of the true parameter values 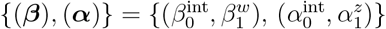.

| Truth $\{(\beta), (\alpha)\}$ | Metric | $\hat{\alpha}_0$ | $\hat{\alpha}_1$ | $\hat{\beta}_0$ | $\hat{\beta}_1$ |
| --- | --- | --- | --- | --- | --- |
| $\{(-1, 0.5), (1, 1)\}$ | avgBias | 0.005 | 0.003 | -0.010 | 0.002 |
|  | avgSEE | 0.035 | 0.033 | 0.086 | 0.098 |
|  | avgSEM | 0.035 | 0.036 | 0.086 | 0.098 |
|  | avgCR (%) | 96.0 | 97.0 | 97.5 | 93.5 |
| $\{(-1, 1), (1, 1)\}$ | avgBias | 0.005 | 0.003 | -0.014 | 0.009 |
|  | avgSEE | 0.035 | 0.036 | 0.110 | 0.124 |
|  | avgSEM | 0.036 | 0.039 | 0.104 | 0.118 |
|  | avgCR (%) | 96.5 | 97.0 | 93.5 | 93.0 |
| $\{(-1, 1), (1, 0.5)\}$ | avgBias | 0.002 | 0.007 | -0.015 | 0.012 |
|  | avgSEE | 0.031 | 0.033 | 0.110 | 0.132 |
|  | avgSEM | 0.033 | 0.034 | 0.108 | 0.122 |
|  | avgCR (%) | 97.0 | 95.0 | 95.5 | 93.5 |

The results are summarized in Table S2.2. Although the total sample size is kept the same as in Simulation Scenario 1, incorporating male data leads to noticeable improvements in model performance, with reductions in both bias and standard errors. In particular, the standard errors for the estimates of ***α*** are now smaller than those for ***β***, indicating that Assumption 1 can substantially enhance estimation efficiency when it holds.

#### S2.2 Simulation for data like those from the Illumina EPIC array

##### S2.2.1 Simulation Scenario 1

In this section, we apply the LVE-X model to data that is assumed to be generated under an Illumina-based methylation framework. Unlike bisulfite sequencing, which yields discrete read counts, Illumina platforms provide methylation proportions that are continuous and bounded between 0 and 1. Accordingly, we adopt the following data-generating mechanism. As before, each cell is assumed to belong to one of three underlying methylation classes, and the simulated methylation proportion for each participant is drawn from a Beta distribution, with its mean specified by the function in Equation (1) in the main Methods Section; the dispersion parameter is fixed at 10. Consistent with the bisulfite sequencing simulations, we use the same three sets of true coefficient values and set the sample size to 200. Model performance is summarized using the same criteria as in the bisulfite sequencing simulation study.

**Table S2.3.** Finite-sample performance in Simulation Scenario 1 (female-only Illumina sequencing–based model) under three configurations of the true parameter values 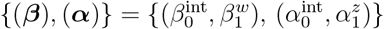

| Truth $\{(\beta), (\alpha)\}$ | Metric | $\hat{\alpha}_0$ | $\hat{\alpha}_1$ | $\hat{\beta}_0$ | $\hat{\beta}_1$ |
| --- | --- | --- | --- | --- | --- |
| $\{(-1, 0.5), (1, 1)\}$ | avgBias | 0.021 | 0.022 | -0.012 | 0.006 |
|  | avgSEE | 0.254 | 0.148 | 0.092 | 0.046 |
|  | avgSEM | 0.249 | 0.148 | 0.088 | 0.048 |
|  | avgCR (%) | 97.0 | 94.5 | 92.5 | 97.0 |
| $\{(-1, 1), (1, 1)\}$ | avgBias | -0.010 | -0.007 | -0.016 | 0.010 |
|  | avgSEE | 0.162 | 0.104 | 0.069 | 0.059 |
|  | avgSEM | 0.171 | 0.112 | 0.074 | 0.063 |
|  | avgCR (%) | 96.5 | 94.5 | 96.5 | 96.5 |
| $\{(-1, 1), (1, 0.5)\}$ | avgBias | 0.044 | 0.020 | -0.002 | -0.004 |
|  | avgSEE | 0.222 | 0.094 | 0.104 | 0.087 |
|  | avgSEM | 0.212 | 0.097 | 0.095 | 0.078 |
|  | avgCR (%) | 94.5 | 96.0 | 90.0 | 90.0 |

From the results presented in Table S2.3, we observe that the model performs well overall according to the reported performance metrics.

##### S2.2.2 Simulation Scenario 2

In this scenario, we also incorporate male data into the modeling framework. We generate methylation data for 100 female participants using the same Illumina-based mechanism described in Section S2.2.1. In addition, we generate data for 100 male participants from a Beta distribution with mean determined solely by *v*(***z***_*i*_) and with the dispersion parameter again fixed at 10, consistent with Section S2.2.1. The same set of performance criteria is used to evaluate the model.

**Table S2.4.** Finite-sample performance in Simulation Scenario 2 (Illumina sequencing–based model incorporating both female and male participants) under three configurations of the true parameter values 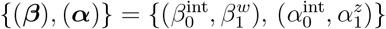

| Truth $\{(\beta), (\alpha)\}$ | Metric | $\hat{\alpha}_0$ | $\hat{\alpha}_1$ | $\hat{\beta}_0$ | $\hat{\beta}_1$ |
| --- | --- | --- | --- | --- | --- |
| $\{(-1, 0.5), (1, 1)\}$ | avgBias | 0.008 | -0.006 | 0.003 | -0.006 |
|  | avgSEE | 0.078 | 0.073 | 0.113 | 0.104 |
|  | avgSEM | 0.070 | 0.074 | 0.099 | 0.099 |
|  | avgCR (%) | 91.5 | 93.0 | 91.5 | 95.0 |
| $\{(-1, 1), (1, 1)\}$ | avgBias | 0.010 | 0.006 | -0.024 | 0.016 |
|  | avgSEE | 0.080 | 0.068 | 0.109 | 0.122 |
|  | avgSEM | 0.070 | 0.073 | 0.111 | 0.123 |
|  | avgCR (%) | 90.5 | 96.0 | 95.5 | 95.0 |
| $\{(-1, 1), (1, 0.5)\}$ | avgBias | 0.006 | 0.003 | -0.014 | 0.019 |
|  | avgSEE | 0.072 | 0.070 | 0.113 | 0.133 |
|  | avgSEM | 0.067 | 0.066 | 0.114 | 0.127 |
|  | avgCR (%) | 93.5 | 94.0 | 96.5 | 92.0 |

From the results shown in Table S2.4, we find that incorporating male data under Assumption 1 improves estimation of ***α***, yielding reduced bias and smaller variance. This pattern is consistent with the gains observed for the bisulfite sequencing–based model.

### S3 Post-hoc Power Calculation for the 2017 RA Dataset

In this section, we present the post-hoc power analysis for the 2017 RA dataset. The objective is to estimate the probability that the 2017 analysis would have successfully detected an effect at a pre-specified significance level (set to 0.05 in our case), assuming that the true effect size corresponds to the one estimated from the 2019 dataset, but with read depths observed in the 2017 study.

The power calculation procedure for each CpG site proceeds as follows:

- For the CpG sites of interest that are present in both the 2017 and 2019 datasets, we extract the estimated regression coefficients from the 2019 analysis.
- Using these coefficients as the assumed true effect sizes, we simulate methylation read counts conditional on the read depths and covariate values observed in the 2017 dataset.
- We then fit the same association model to the simulated 2017 data and extract the *p*-value corresponding to the ACPA or RA term.
- The above steps are repeated 2,000 times. The empirical power is estimated as the proportion of simulations in which the *p*-value is less than 0.05.

All estimated post-hoc power for overlap sites has been recorded in the accompanying Excel file.

## Supplemental Figures

**Figure S1.**
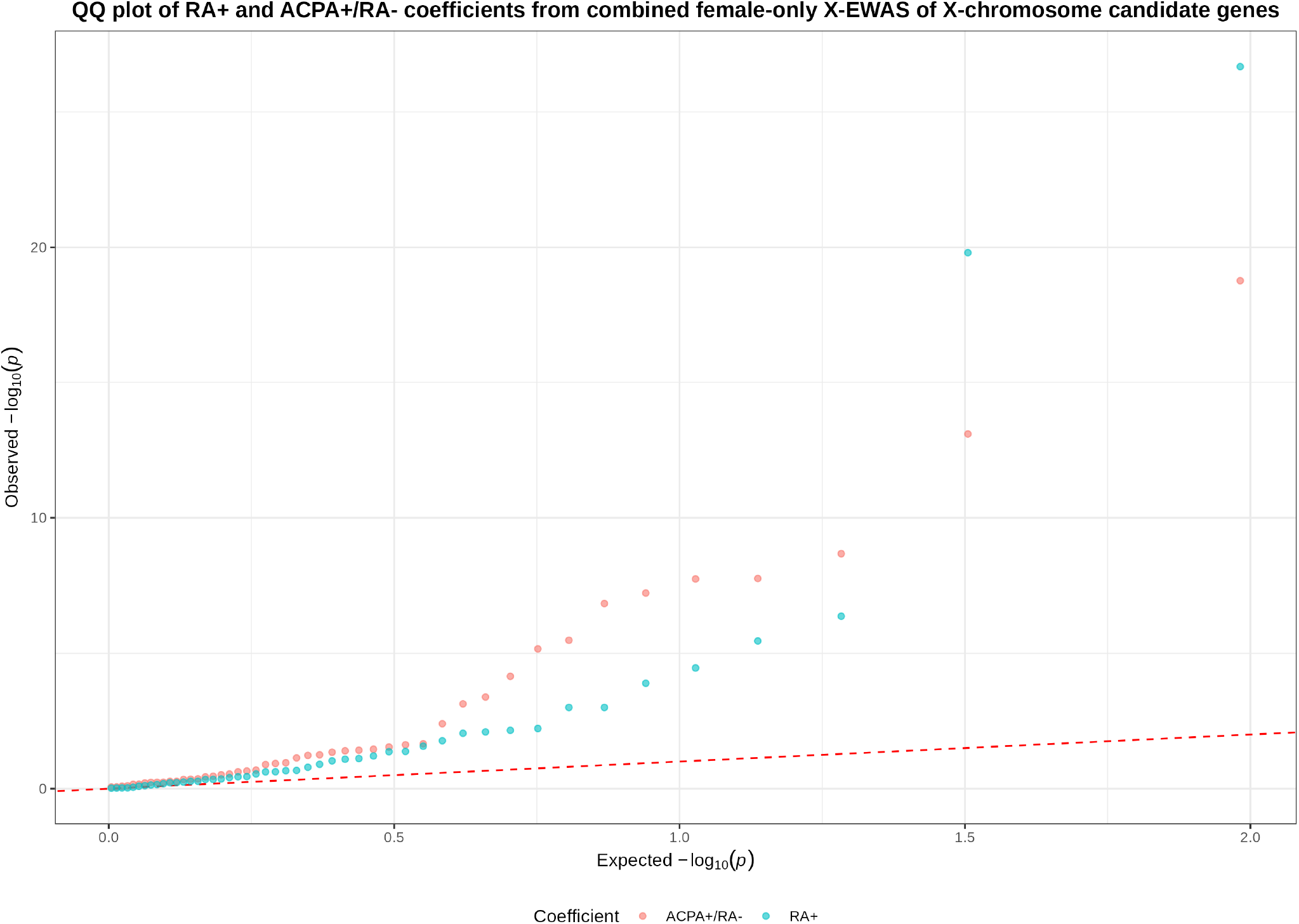
Quantile–quantile (Q–Q) plots comparing the observed and expected *−* log_10_(*p*) values for the ACPA+/RA*−* and RA+ coefficients from the female-only X-AS of candidate genes in the combined dataset, obtained by integrating the 2019 and 2017 cohorts.

**Figure S2.**
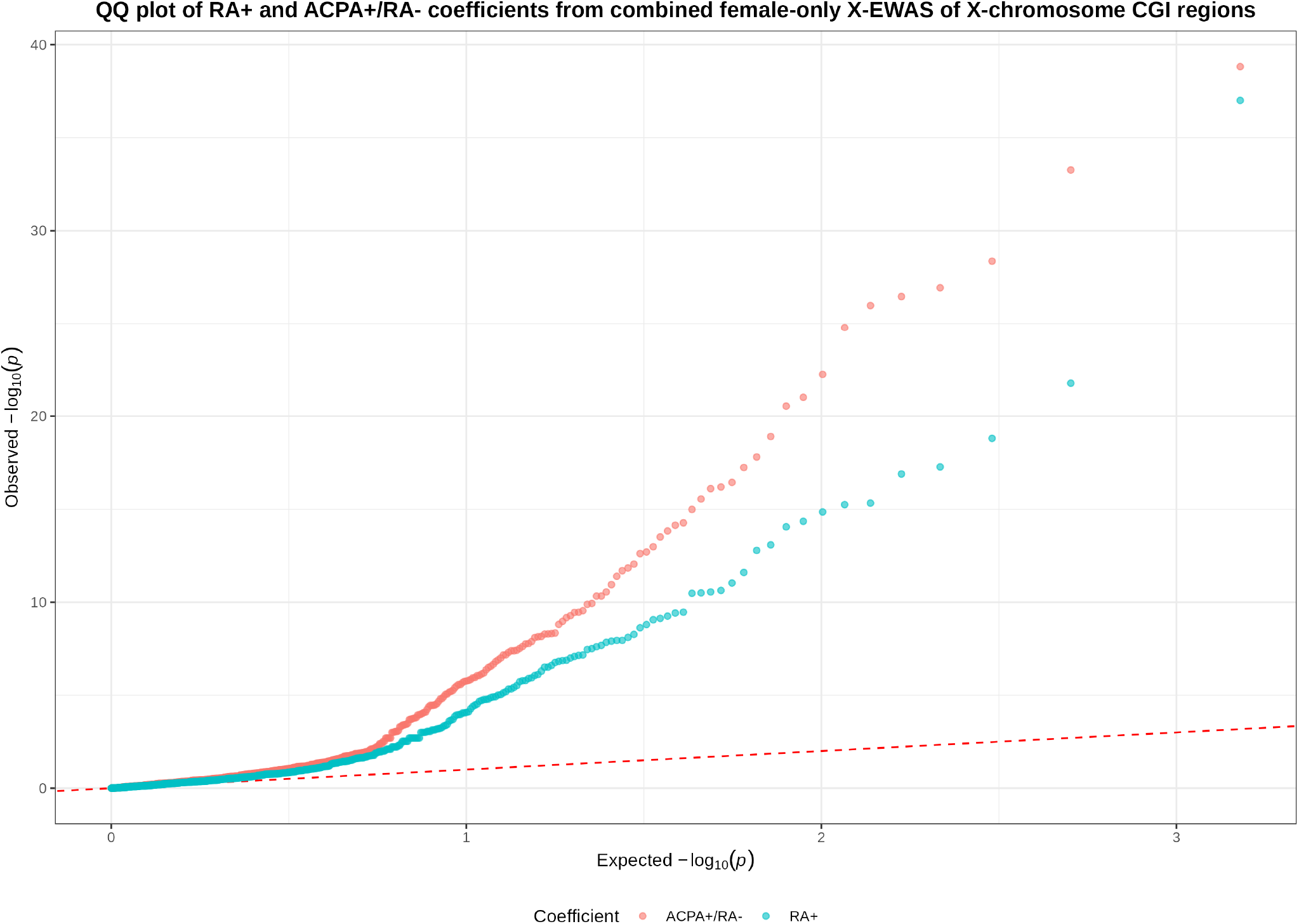
Quantile–quantile (Q–Q) plots comparing the observed and expected *−* log_10_(*p*) values for the ACPA+/RA*−* and RA+ coefficients from the female-only X-AS of CpG islands in the combined dataset, obtained by integrating the 2019 and 2017 cohorts.

**Figure S3.**
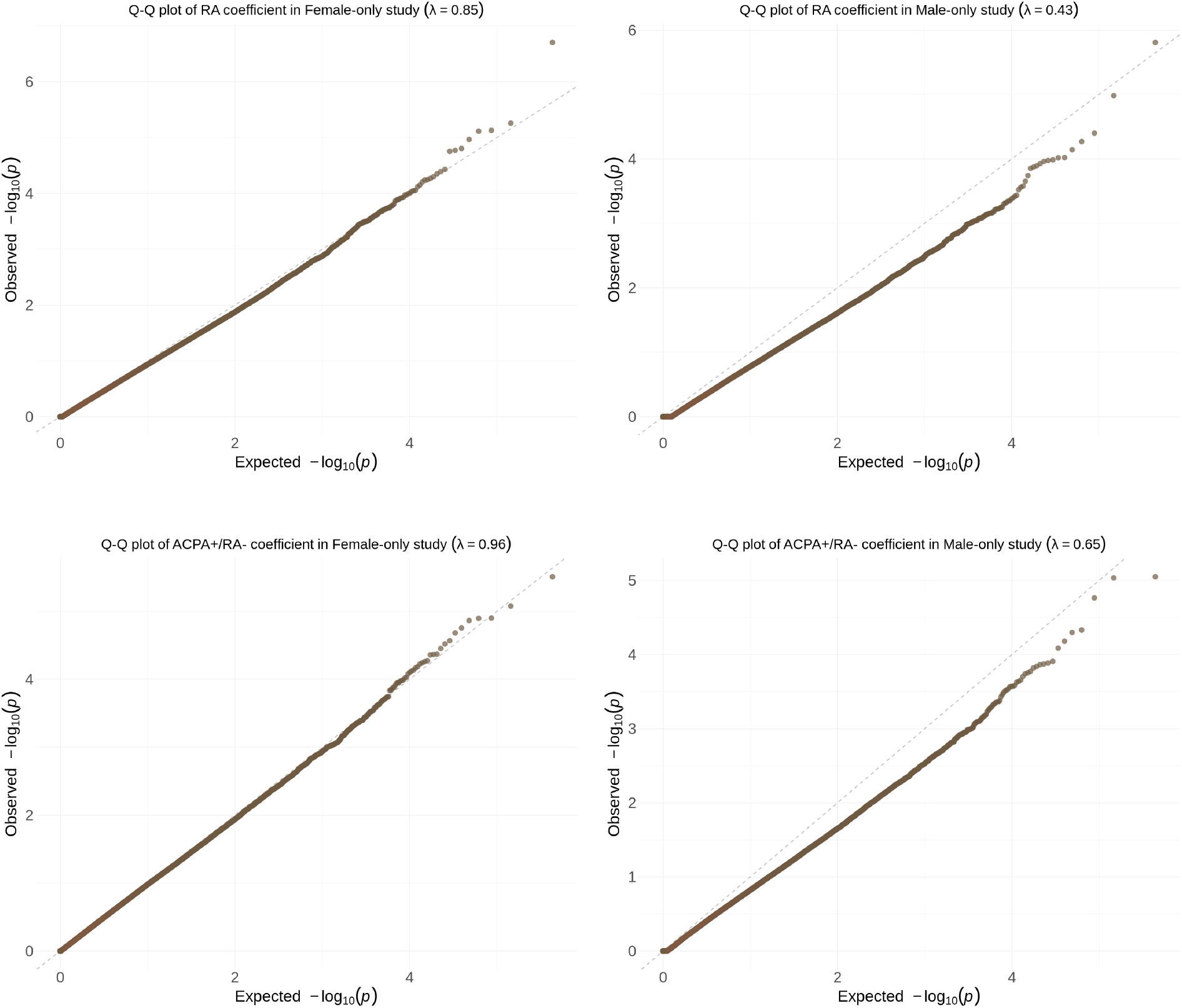
Quantile–quantile (Q–Q) plots comparing observed and expected *−* log_10_(*p*) values from sex-stratified X-AS analyses of single CpGs in the 2019 RA dataset. The top row shows the comparison of RA+ versus ACPA*−*/RA*−* individuals, whereas the bottom row shows the comparison of ACPA+/RA*−* versus ACPA*−*/RA*−* individuals. The left and right panels correspond to females and males, respectively. Here, *λ* denotes the genomic inflation factor.

**Figure S4.**
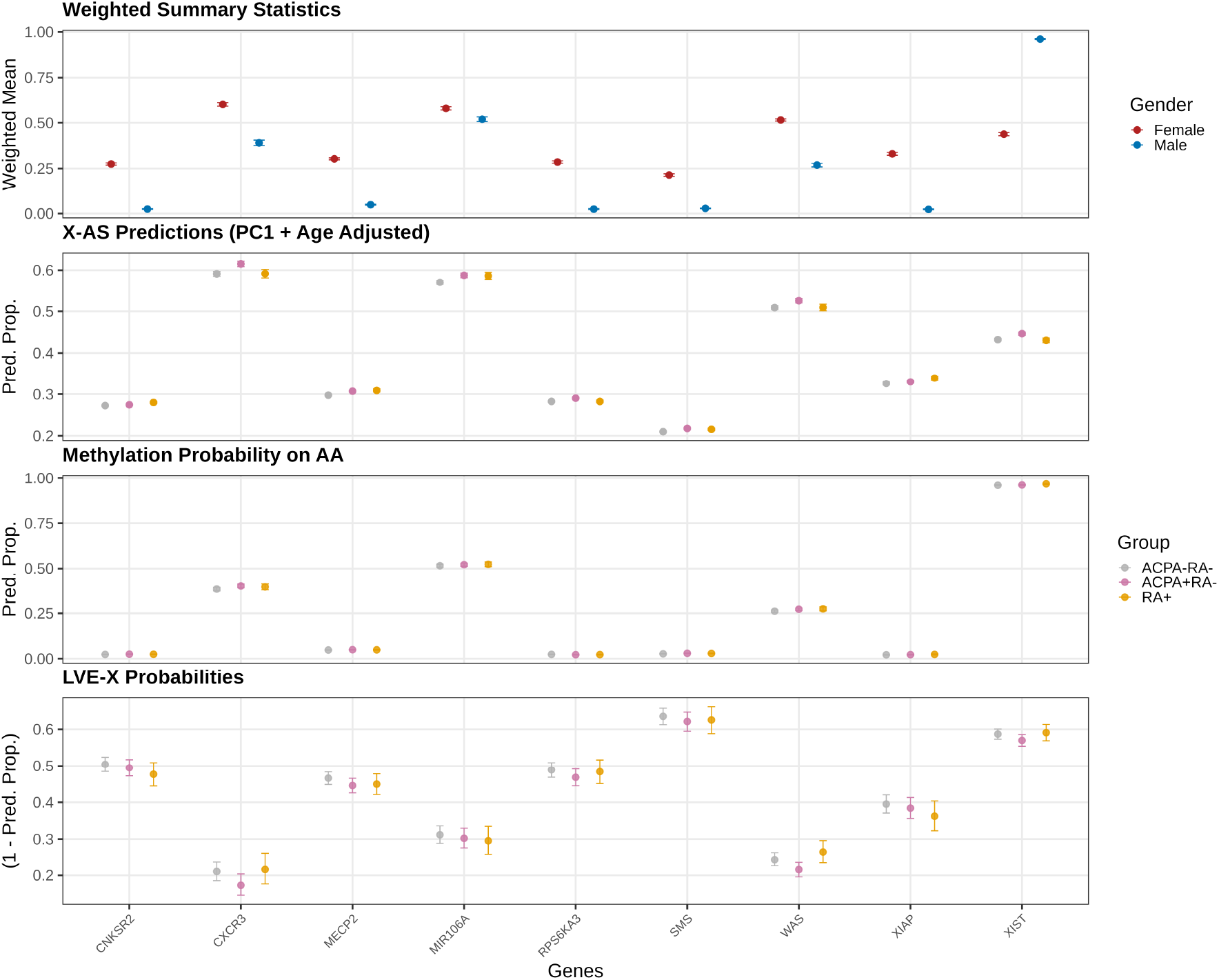
Pipeline and LVE-X model results from aggregate analyses at nine candidate genes on the X chromosome: *CNKSR2, CXCR3, MECP2, MIR106A, RPS6KA3, SMS, WAS, XIAP*, and *XIST*. The top panel shows mean methylation levels weighted by read depths, *±* one standard deviation, for males and females. In the second panel, results of the X-AS analysis in females are shown as predicted mean methylation levels in three phenotypic groups defined by RA and ACPA status. For these predictions, age is set to the mean age, and the first principal component of cell type proportions is set to zero (*±* 95% confidence intervals). In the third panel, the predicted methylation proportions (*v*(.); *±* 95% confidence intervals) are shown for the AA alleles from the LVE-X model, with age set to mean age, and the first principal component of cell type proportions set to zero. In the bottom panel, one minus the predicted methylation proportions (*π*(.); *±* 95% confidence intervals) are shown for the IA alleles from the LVE-X model.

**Figure S5.**
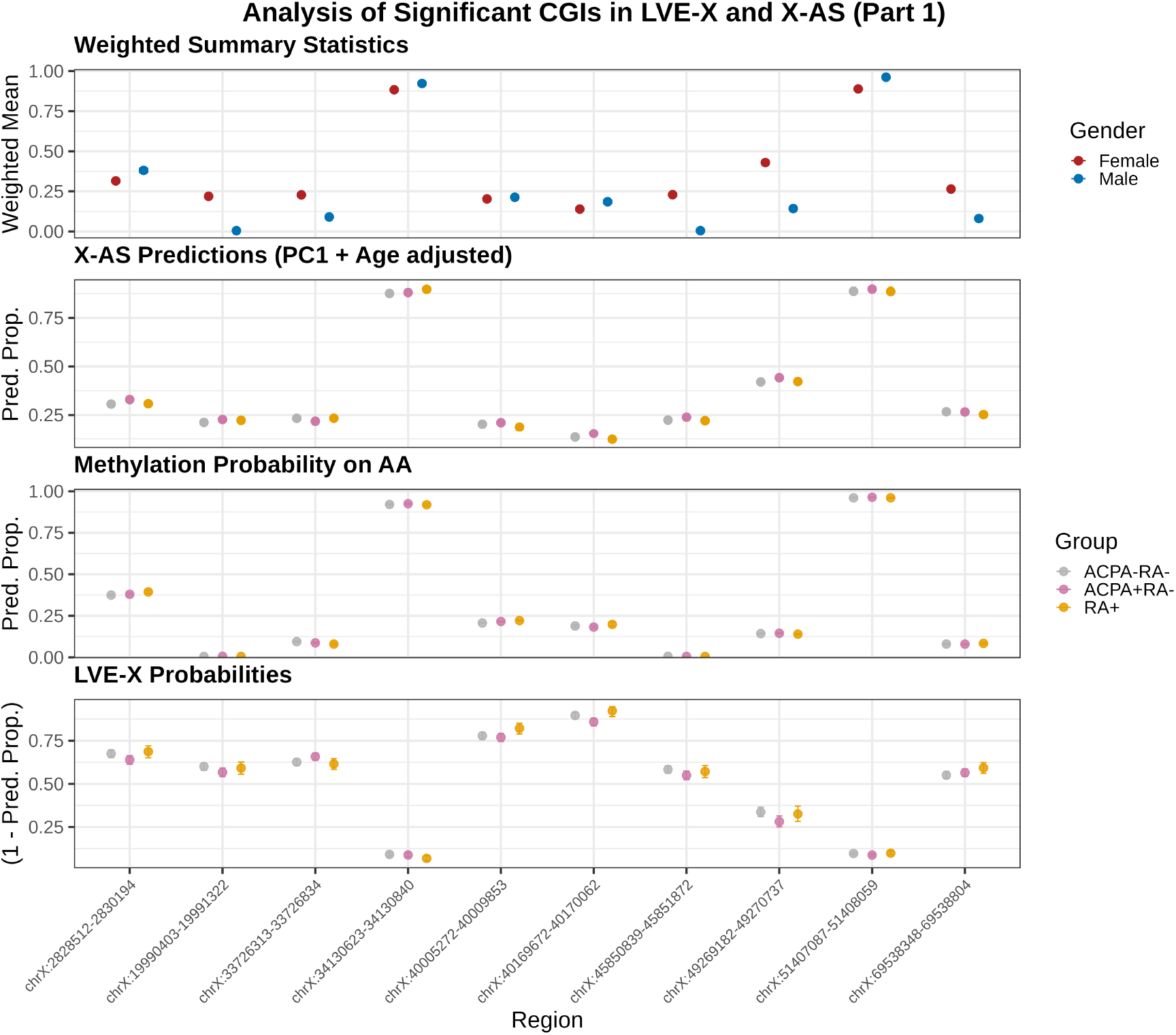
Pipeline and LVE-X model results from aggregate analyses of 10 CpG islands (CGIs) on the X chromosome selected based on significant findings from both LVE-X probability analyses and X-AS. The top panel shows mean methylation levels weighted by read depths, *±* one standard deviation, for males and females. In the second panel, results of the X-AS analysis in females are shown as predicted mean methylation levels in three phenotypic groups defined by RA and ACPA status. For these predictions, age is set to the mean age, and the first principal component of cell type proportions is set to zero (*±* 95% confidence intervals). In the third panel, the predicted methylation proportions (*v*(.); *±* 95% confidence intervals) are shown for the AA alleles from the LVE-X model, with age set to mean age, and the first principal component of cell type proportions set to zero. In the bottom panel, one minus the predicted methylation proportions (*π*(.);*±* 95% confidence intervals) are shown for the IA alleles from the LVE-X model.

**Figure S6.**
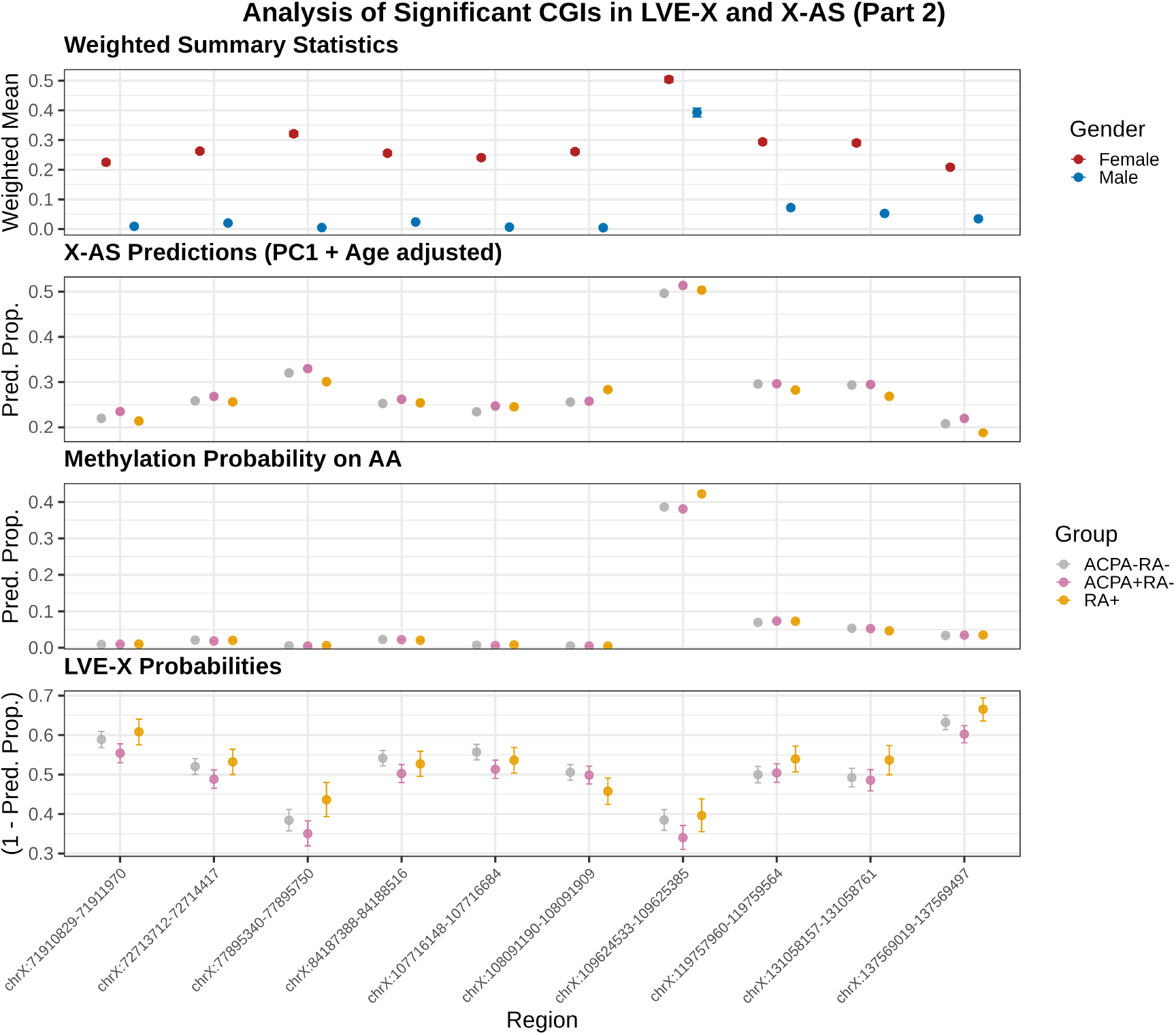
Pipeline and LVE-X model results from aggregate analyses of 10 CpG islands (CGIs) on the X chromosome selected based on significant findings from both LVE-X probability analyses and X-AS. The top panel shows mean methylation levels weighted by read depths, *±* one standard deviation, for males and females. In the second panel, results of the X-AS analysis in females are shown as predicted mean methylation levels in three phenotypic groups defined by RA and ACPA status. For these predictions, age is set to the mean age, and the first principal component of cell type proportions is set to zero (*±* 95% confidence intervals). In the third panel, the predicted methylation proportions (*v*(.); *±* 95% confidence intervals) are shown for the AA alleles from the LVE-X model, with age set to mean age, and the first principal component of cell type proportions set to zero. In the bottom panel, one minus the predicted methylation proportions (*π*(.);*±* 95% confidence intervals) are shown for the IA alleles from the LVE-X model.

**Figure S7.**
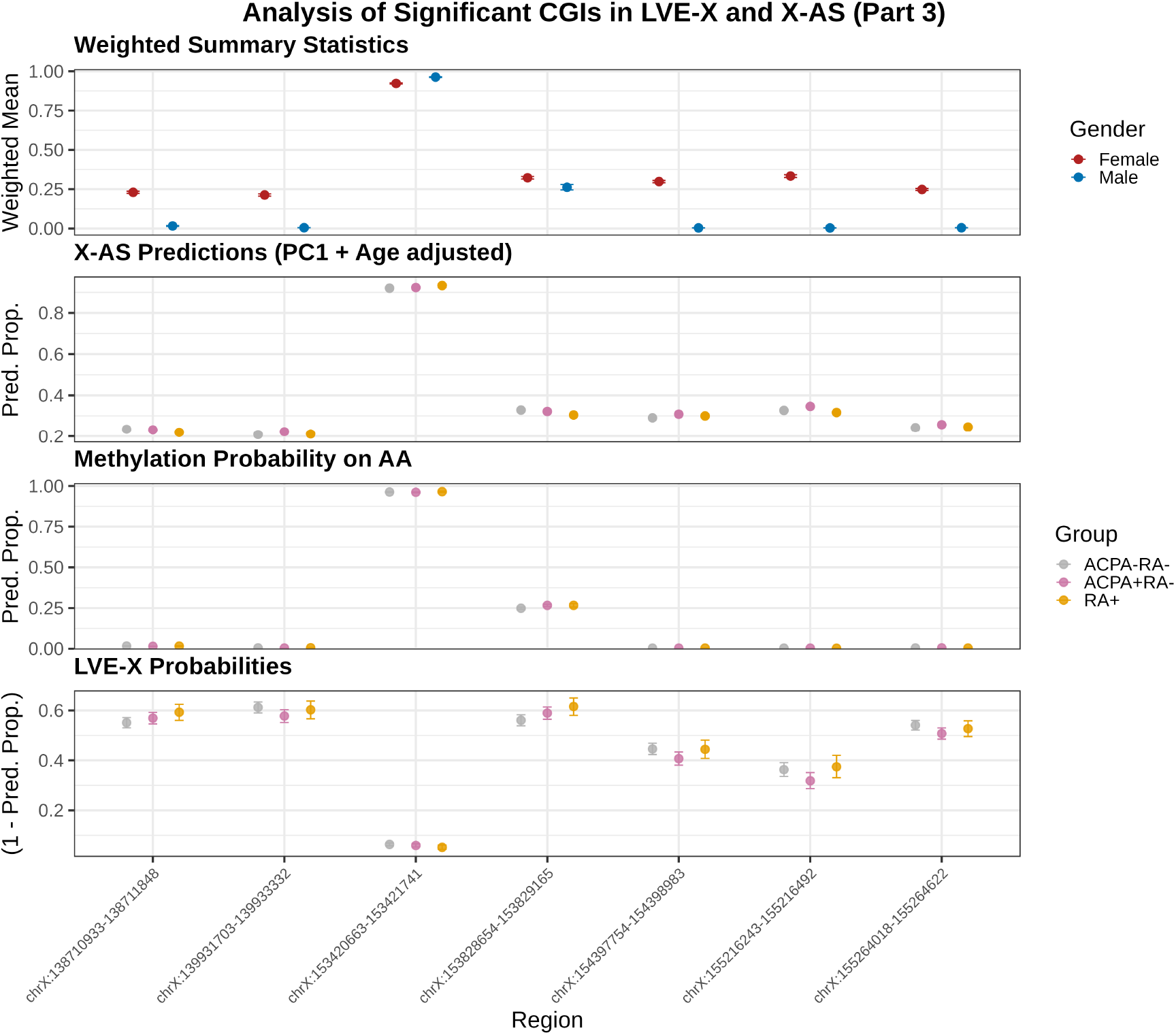
Pipeline and LVE-X model results from aggregate analyses of 7 CpG islands (CGIs) on the X chromosome selected based on significant findings from both LVE-X probability analyses and X-AS. The top panel shows mean methylation levels weighted by read depths, *±* one standard deviation, for males and females. In the second panel, results of the X-AS analysis in females are shown as predicted mean methylation levels in three phenotypic groups defined by RA and ACPA status. For these predictions, age is set to the mean age, and the first principal component of cell type proportions is set to zero (*±* 95% confidence intervals). In the third panel, the predicted methylation proportions (*v*(.); *±* 95% confidence intervals) are shown for the AA alleles from the LVE-X model, with age set to mean age, and the first principal component of cell type proportions set to zero. In the bottom panel, one minus the predicted methylation proportions (*π*(.);*±* 95% confidence intervals) are shown for the IA alleles from the LVE-X model.

**Figure S8.**
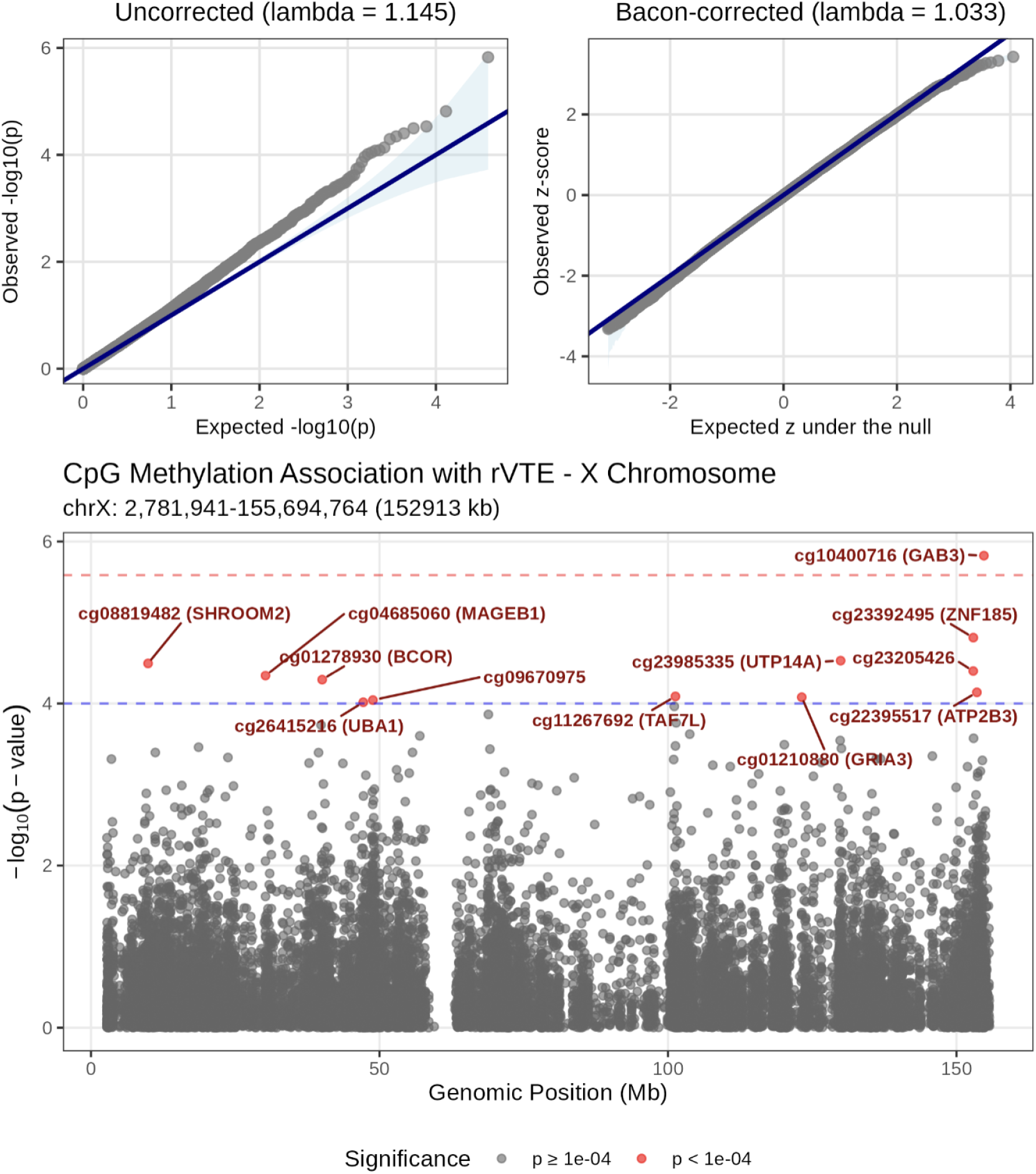
QQ plots and X-chromosome regional association plot for the female X-AS of rVTE status. QQ plots display the distribution of observed versus expected *−* log_10_(*p*) values before bacon correction and observed versus expected Z statistics after bacon correction. The X-chromosome regional association plot shows *−* log_10_(*p*) values for all 19,517 CpG sites across the X chromosome. Horizontal red and blue lines denote the Bonferroni-corrected (*p <* 2.6 *×* 10^*−*6^) and suggestive (*p <* 1 *×* 10^*−*4^) significance thresholds, respectively.

**Figure S9.**
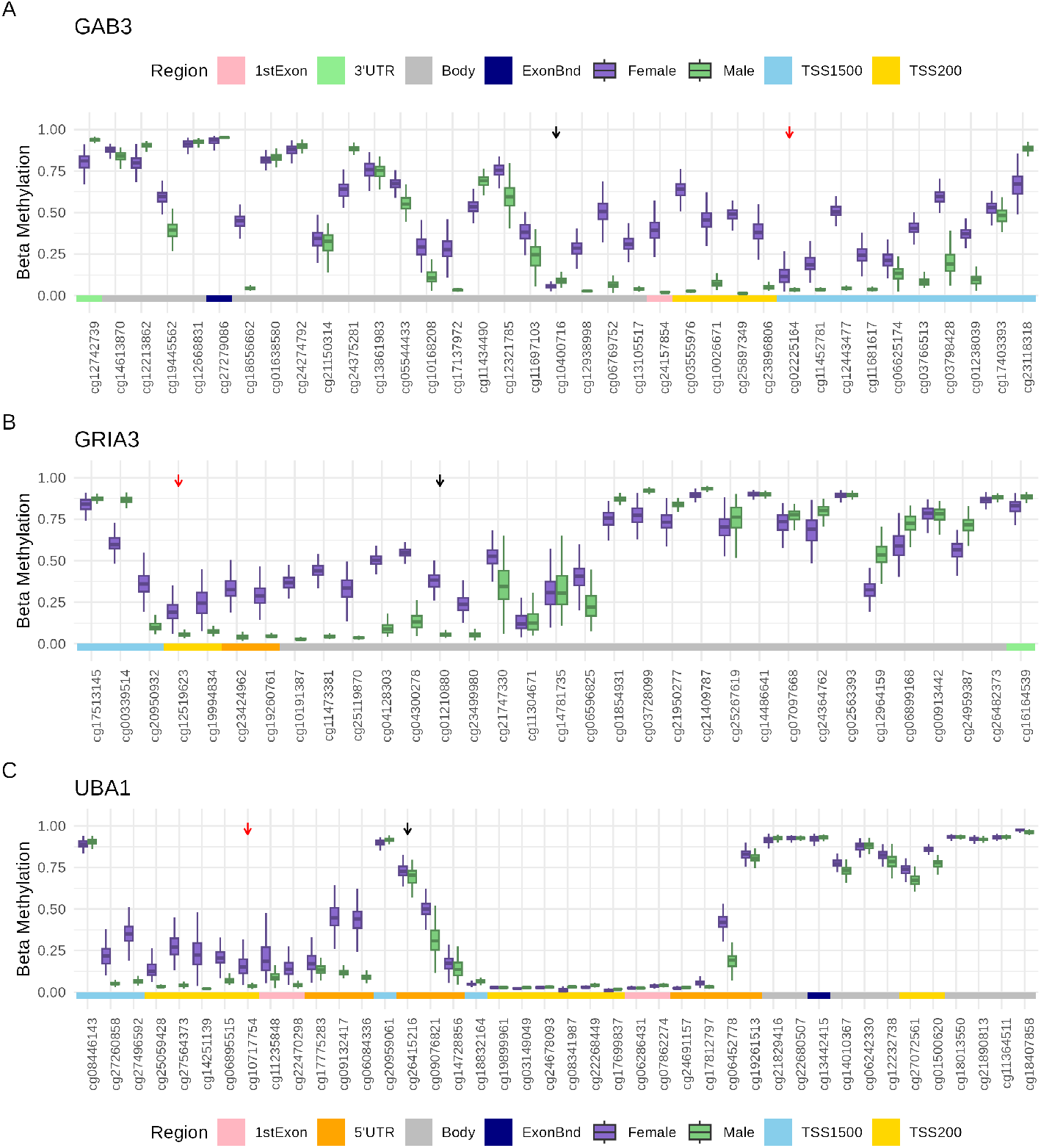
DNA methylation levels of probes across the rVTE-associated gene, stratified by sex. Probes on the x-axis are ordered by genomic position, and colored regions indicate genomic annotations based on the Illumina EPIC manifest (GENCODE v12). Females are shown in purple and males in green. Black arrows indicate the rVTE-associated CpG, and red arrows highlight CpGs showing sex-differential methylation within 1–10 kb of the associated site. A) *GAB3* ; B) *GRIA3* ; and C) UBA1.

## Supplemental Tables

**Table S1.**
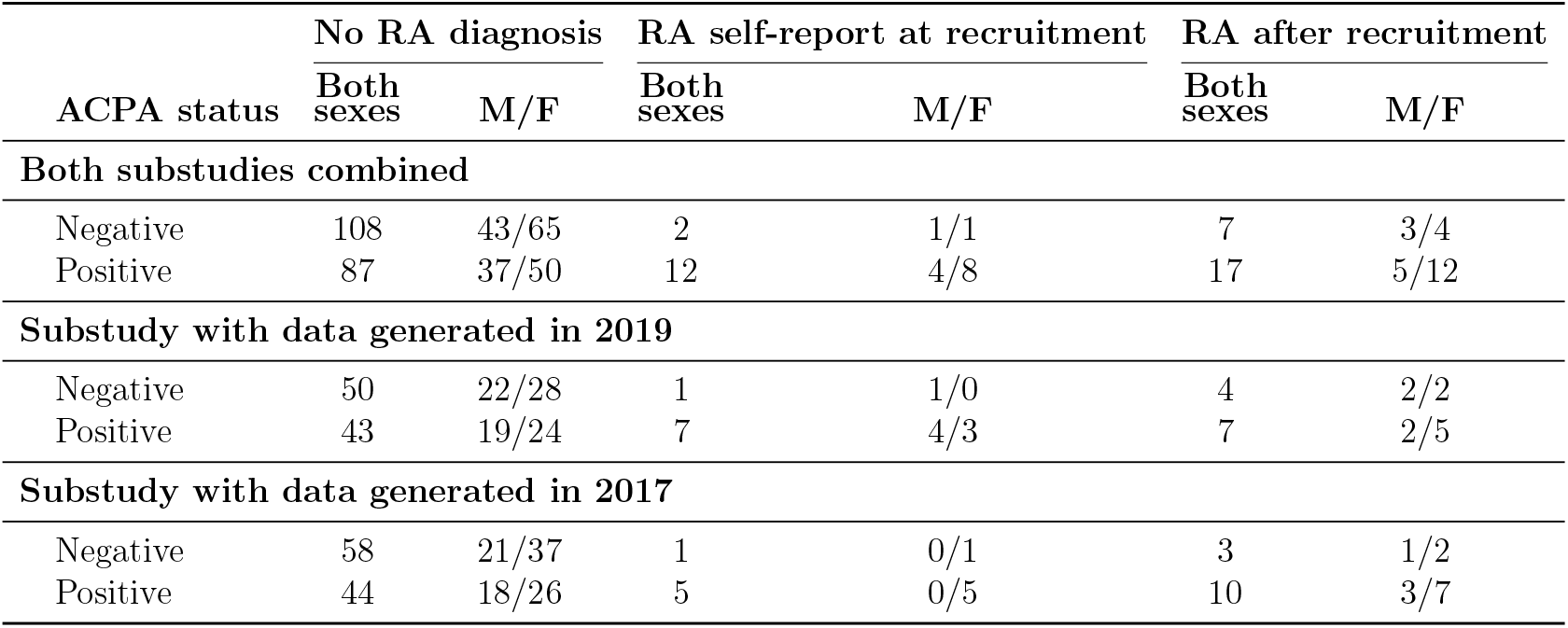
Participants in the two substudies of DNA methylation and ACPA positivity, and their RA status. M/F indicate numbers by sex (Male/Female).

**Table S2.** Demographic information and RA diagnosis status for the two RA datasets.

|  | 2017 data |  | 2019 data |  |
| --- | --- | --- | --- | --- |
|  | Mean (SD) / *n* (%) | Range | Mean (SD) / *n* (%) | Range |
| Age | 54.751 (7.726) | [40.350, 69.940] | 54.181 (7.866) | [40.740, 70.220] |
| RA diagnosis status, *n* (%) |  |  |  |  |
| No RA diagnosis | 102 (84.3) | - | 93 (83.0) | - |
| RA self-report at recruitment | 6 (5.0) | - | 8 (7.1) | - |
| RA after recruitment | 13 (10.7) | - | 11 (9.8) | - |
| Smoker, *n* (%) |  |  |  |  |
| Current | 26 (21.5) | - | 18 (16.1) | - |
| Past | 47 (38.8) | - | 48 (42.9) | - |
| Never | 4 (3.3) | - | 4 (3.6) | - |
| Missing | 44 (36.4) | - | 42 (37.5) | - |
| Blood cell proportions |  |  |  |  |
| Monocyte | 0.077 (0.022) | [0.037, 0.186] | 0.079 (0.019) | [0.037, 0.14] |
| Lymphocyte | 0.280 (0.068) | [0.131, 0.504] | 0.286 (0.072) | [0.110, 0.420] |
| Neutrophil | 0.613 (0.078) | [0.392, 0.783] | 0.604 (0.082) | [0.408, 0.81] |
| Eosinophil | 0.023 (0.015) | [0.004, 0.120] | 0.025 (0.017) | [0.002, 0.121] |
| Basophil | 0.007 (0.004) | [0.000, 0.023] | 0.006 (0.004) | [0, 0.023] |
| PC1 of cell type proportions | $3.73 \times 10^{-17}$ (0.102) | [-0.224, 0.306] | $4.17 \times 10^{-17}$ (0.108) | [-0.272, 0.236] |

**Table S3.** CpG coverage under a range of filtering criteria in the 2017 and 2019 RA datasets

| **A. CpG sites retained under several filters for read coverage** |  |  |  |
| --- | --- | --- | --- |
| Filter criterion | 2017 RA dataset | 2019 RA dataset | Overlap |
| Original sites | 1,245,869 | 1,229,858 | 1,214,993 |
| After QC | 241,127 | 192,834 | 175,259 |
| > 2 reads in > 20 samples | 241,149 | 192,832 | 175,280 |
| ≥ 5 reads in ≥ 1 sample | 267,564 | 423,191 | 237,125 |
| ≥ 5 reads in ≥ 50% samples | 121,246 | 158,053 | 117,311 |
| ≥ 5 reads in ≥ 80% samples | 81,875 | 133,776 | 78,898 |
| ≥ 10 reads in ≥ 50% samples | 63,484 | 123,744 | 60,779 |
| **B. CpG sites mostly methylated or unmethylated in > 80% individuals** |  |  |  |
| Criterion | Subgroup | 2017 RA dataset | 2019 RA dataset |
| Methylation rate < 20% | All Participants | 40,665 | 38,950 |
| Methylation rate < 20% | Males | 98,529 | 95,481 |
| Methylation rate < 20% | Females | 36,852 | 38,763 |
| Methylation rate ≥ 70% | All Participants | 662,870 | 694,848 |
| Methylation rate ≥ 70% | Males | 763,317 | 810,854 |
| Methylation rate ≥ 70% | Females | 607,217 | 618,396 |

**Table S4.**
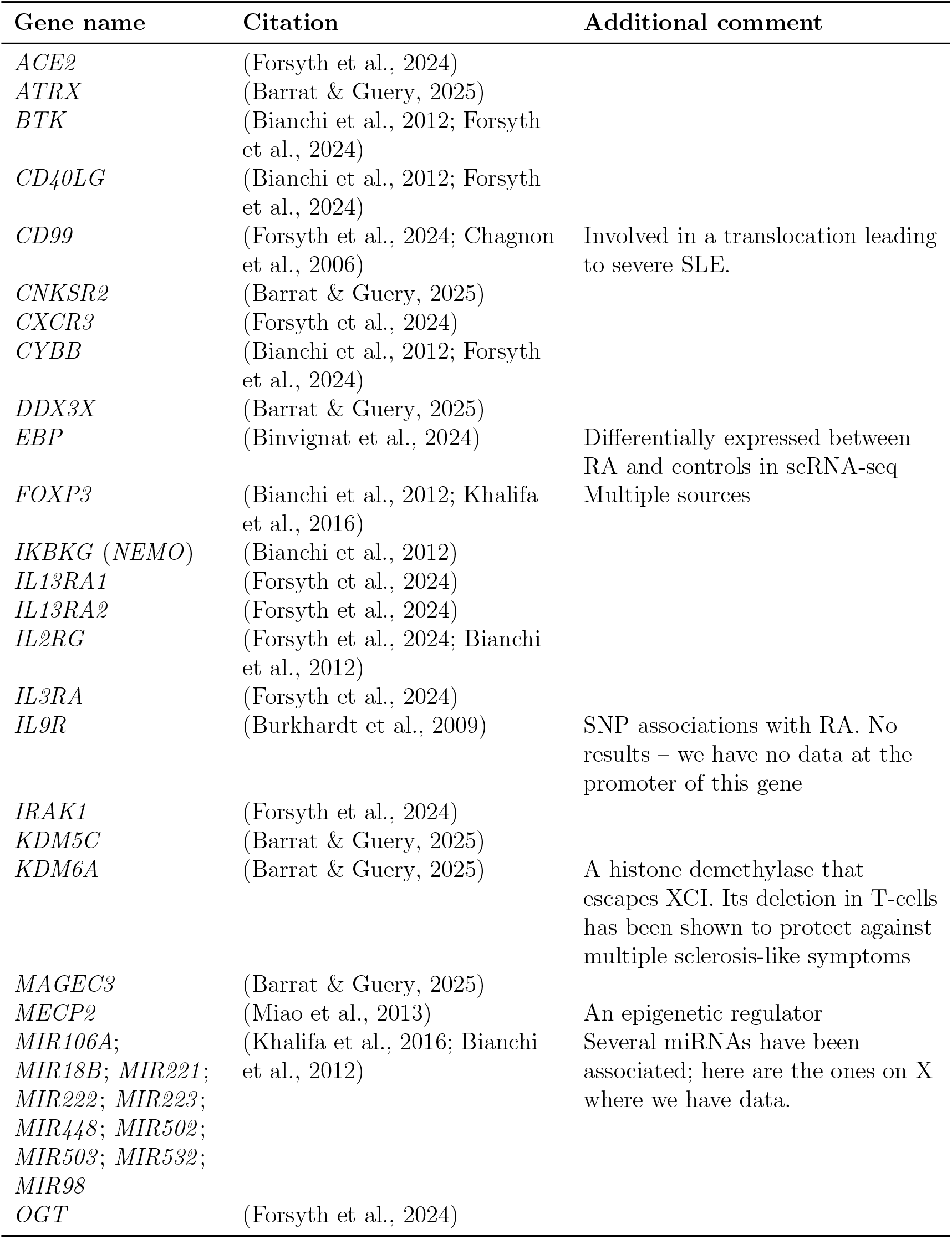
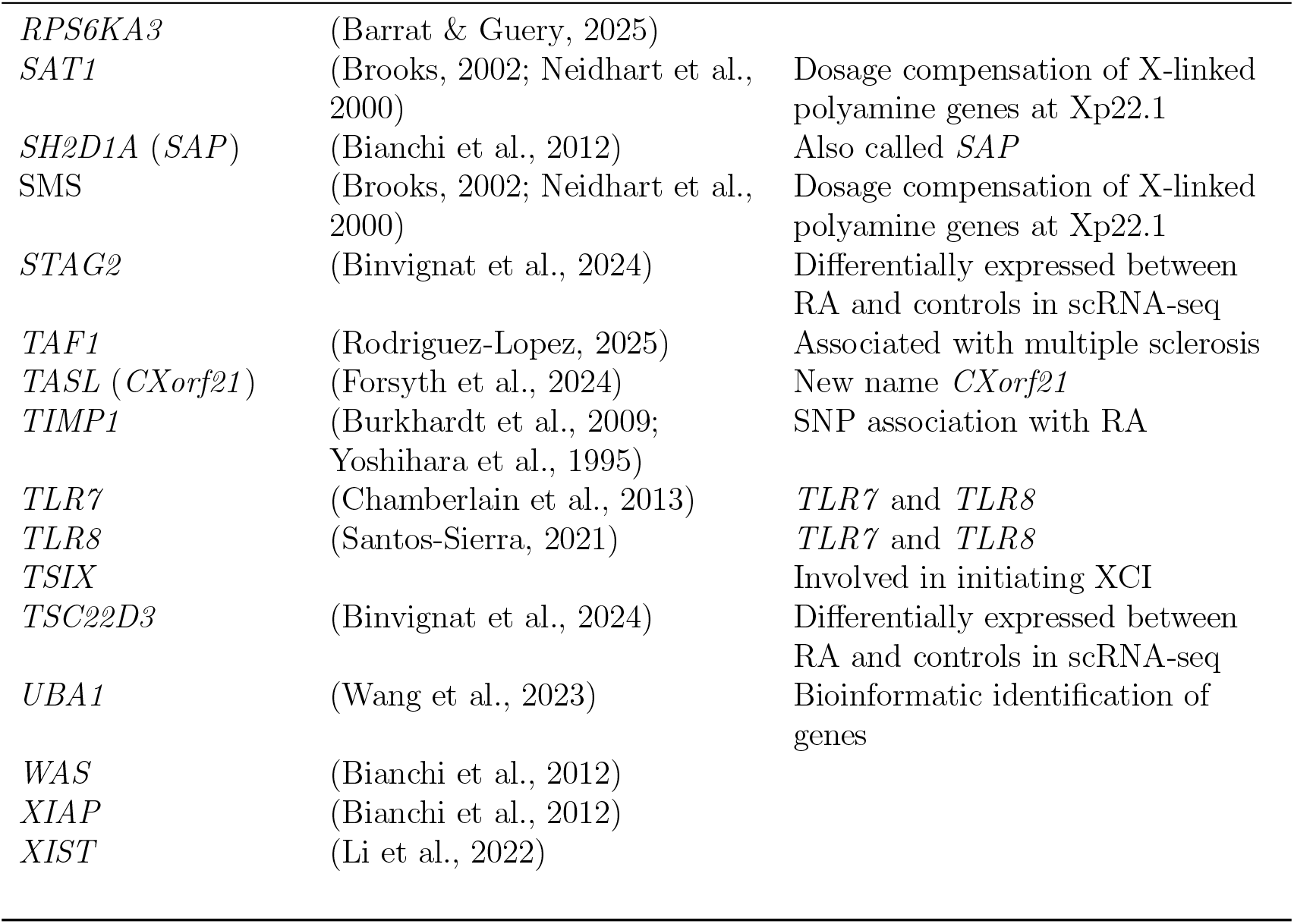
References and additional comments for the selected candidate genes.

**Table S5.** Demographic information of the rVTE dataset.

|  | Female |  |  |  | Male |  |  |  |
| --- | --- | --- | --- | --- | --- | --- | --- | --- |
|  | VTE (n=175) |  | rVTE (n=32) |  | VTE (n=141) |  | rVTE (n=69) |  |
|  | Mean (SD) | Range | Mean (SD) | Range | Mean (SD) | Range | Mean (SD) | Range |
| Age | 51.3 (19.4) | [18.0–93.5] | 57 (20.6) | [20.2–86.9] | 56.3 (15.4) | [22.2–90.3] | 52.8 (14.8) | [18.6–84] |
| BMI<br>(kg/m<sup>2</sup>) | 28.6 (8.0) | [14.8–78] | 32.5 (9.6) | [19.1–52.3] | 29.5 (6) | [18.3–71.1] | 29.3 (7.3) | [17.9–60.1] |
| Time to<br>recurrence | 6.3 (2.8) | [0.07–9.2] | 2.5 (2.4) | [0.02–7.4] | 5.6 (3) | [0.02–9.77] | 2.3 (2.3) | [0.04–8.09] |

